# Near-native state imaging by cryo-soft-X-ray tomography reveals remodelling of multiple cellular organelles during HSV-1 infection

**DOI:** 10.1101/2021.10.11.463900

**Authors:** Kamal L. Nahas, Viv Connor, Katharina M. Scherer, Clemens F. Kaminski, Maria Harkiolaki, Colin M. Crump, Stephen C. Graham

## Abstract

Herpes simplex virus-1 (HSV-1) is a large, enveloped DNA virus and its assembly in the cell is a complex multi-step process during which viral particles interact with numerous cellular compartments such as the nucleus and organelles of the secretory pathway. Transmission electron microscopy and fluorescence microscopy are commonly used to study HSV-1 infection. However, 2D imaging limits our understanding of the 3D geometric changes to cellular compartments that accompany infection and sample processing can introduce morphological artefacts that complicate interpretation. In this study, we used soft X-ray tomography to observe differences in whole-cell architecture between HSV-1 infected and uninfected cells. To protect the near-native structure of cellular compartments we used a non-disruptive sample preparation technique involving rapid cryopreservation, and a fluorescent reporter virus was used to facilitate correlation of structural changes with the stage of infection in individual cells. We observed viral capsids and assembly intermediates interacting with nuclear and cytoplasmic membranes. Additionally, we observed differences in the morphology of specific organelles between uninfected and infected cells. The local concentration of cytoplasmic vesicles at the juxtanuclear compartment increased and their mean width decreased as infection proceeded, and lipid droplets transiently increased in size. Furthermore, mitochondria in infected cells were elongated and highly branched, suggesting that HSV-1 infection alters the dynamics of mitochondrial fission/fusion. Our results demonstrate that high-resolution 3D images of cellular compartments can be captured in a near-native state using soft X-ray tomography and have revealed that infection causes striking changes to the morphology of intracellular organelles.

**Importance:** Ultrastructural changes to the morphology and organization of cellular compartments during herpes simplex virus-1 (HSV-1) infection have not previously been studied under near-physiological conditions. In this study, soft X-ray tomography was used to image the ultrastructure of vitrified cells during HSV-1 infection, identifying striking changes to the abundance and organization of cytoplasmic vesicles and mitochondria. The concentration of vesicles in the juxtanuclear region increased with time post infection, which could represent an increasing supply of vesicles to support capsid envelopment, and there is a transient increase in the size of lipid droplets in infected cells. Furthermore, we show that mitochondria elongate and form highly-branched networks as infection progresses. These findings offer insight into stages of virion morphogenesis and the cellular response to infection, highlighting the utility of cryo-soft-X-ray tomography for monitoring the near-native state ultrastructure of infected cells.

## Introduction

Herpes simplex virus-1 (HSV-1) is a large, enveloped DNA virus in the *Alphaherpesvirinae* subfamily of *Herpesviridae* that establishes a persistent life-long latent infection in sensory and sympathetic neurons, occasionally reactivating to cause lytic replication in oral or genital mucosal epithelial cells that culminates in cold sores and genital herpes, respectively^1^. The production of viral particles during lytic replication is a complex process involving multiple cellular compartments^2–6^.

In the first step of virion morphogenesis, capsid assembly and genome packaging occur in the nucleus^7^. Fully formed nucleocapsids must cross the nuclear envelope to migrate into the cytoplasm to undergo the latter stages of virus assembly – a process that involves close interaction between nucleocapsids and the membranes of the nuclear envelope. Unlike individual proteins, the nucleocapsids are too large to pass through nuclear pores and must therefore first bud into the perinuclear space through the inner-nuclear membrane, forming a perinuclear viral particle (primary envelopment). The envelope of this particle subsequently fuses with the outer-nuclear membrane to release the nucleocapsid into the cytoplasm (de-envelopment)^8–11^. Numerous copies of multiple (≥ 23) nuclear and cytoplasmic viral proteins deposit on their surface of nucleocapsids, forming the amorphous proteinaceous layer known as the tegument^12^. Tegument proteins have multiple important roles during infection, including the promotion of virion maturation^2, 3, 6^. Several cytoplasmic compartments are essential to virion morphogenesis: viral proteins are synthesized and modified in the endoplasmic reticulum and Golgi complex and, in a process known as secondary envelopment, nucleocapsids acquire their membrane envelope from cytoplasmic vesicles that are thought to be derived from the *trans*-Golgi network and the endosomal system^2, 3, 13^. In addition to compartments directly involved in virion assembly, the cytoskeleton and other cellular organelles, such as mitochondria and lysosomes, can become remodelled in response to infection^14–16^. Understanding how the morphology and organization of cellular compartments change during infection could illuminate their involvement in virion morphogenesis and in the cellular response to HSV-1 infection.

Previous studies to characterize remodelling of cellular compartments have identified numerous changes that accompany HSV-1 infection, including the fragmentation of the Golgi complex and the condensation of the endoplasmic reticulum around the nuclear rim^17, 18^. A more comprehensive study has recently been carried out using a recombinant form of HSV-1, known as the “timestamp” reporter virus, expressing fluorescent chimeras of the early protein ICP0 and the late protein gC to distinguish between early and late stages of infection^16^. Eight cellular compartments were compared between uninfected and timestamp virus-infected human TERT-immortalized human foreskin fibroblast (HFF-hTERT) cells, with high-resolution spatial data collected using structured illumination microscopy (SIM) and expansion microscopy. Numerous changes in the morphology of cellular compartments were observed as infection progressed, such as fragmentation of the Golgi complex at late stages of infection, concentration of endosomes and lysosomes at a juxtanuclear compartment, and elongation of mitochondria^16^. Mitochondrial morphology is known to vary in response to cellular energy demand, oxidative stress, virus infection, and other stimuli^19–24^. For example, varying energy demand within a cell affects mitochondrial length in order to tune the level of ATP production, and fusion of normal and damaged mitochondria during high oxidative stress dilutes the impact of reactive oxygen species on mitochondrial function^19, 20, 24^. Furthermore, mitochondria associate with the cellular microtubule network, which is known to be altered via breakdown and dispersal of the microtubule organising centre during HSV-1 infection^25^. Deregulation of microtubule dynamics may also affect the organisation of cytoplasmic vesicles and their migration from the perinuclear region towards the cell surface.

The extent to which sample preparation strategies alter the morphology of cellular structures remains poorly understood and it is possible that disruptive techniques such as immunostaining or sample expansion could introduce artefacts in cellular ultrastructure^26–28^. Moreover, it is not clear if the changes to cellular compartments that have been observed previously are consistent across different cell types used to study HSV-1 infection. Soft X-ray tomography of cryopreserved samples (cryoSXT) offers an attractive alternative for the imaging of biological samples in a near-native state. Soft X-rays used for cryoSXT have a lower energy (∼500 eV)^29^ and longer wavelength than the “hard” X-rays typically used for medical imaging (∼15–30 keV)^30^ or X-ray crystallography (∼6–20 keV)^31^. The wavelengths of soft X-rays used for cryoSXT are in the “water window” where carbon-rich structures in the cell such as membranes produce considerable contrast whereas oxygen-rich structures such as the “watery” cytosol remain transparent, thereby enabling cellular compartments to be observed^29^. This label-free technique can be used to image the ultrastructure of infected (and control) cells, monitoring the 3D geometry and organization of cellular compartments^32^. Furthermore, by using non-disruptive cryopreservation protocols, such as plunge cryocooling in the case of cryoSXT, the ultrastructure of samples can be preserved in a near-native state for imaging^33^. CryoSXT is particularly suitable for monitoring mitochondria because these cellular compartments produce a lot of contrast owing to their carbon-rich cristae and matrix proteins^29^.

In this study we applied cryoSXT to the study of ultrastructural changes that accompany HSV-1 infection of human osteosarcoma U2OS cells, allowing comparison with previous fluorescence microscopy investigations of HSV-1–infected HFF-hTERT cells^16^. The stage of HSV-1 infection in each individual cell subjected to cryoSXT interrogation was determined by use of the timestamp HSV-1 reporter virus and fluorescence cryo-microscopy. Although a few differences were observed between the extent of Golgi fragmentation and the subcellular distribution of ICP0, we determined that remodelling of cytoplasmic vesicles and mitochondria during infection was largely similar between these cultured cells. Furthermore, the high resolution afforded by cryoSXT revealed that mitochondria become highly branched during HSV-1 infection and that lipid droplets are enlarged at early times post-infection.

## Results

### HSV-1 viral particles and assembly intermediates are detectable by cryoSXT

Transmission electron microscopy (TEM) has been used extensively to visualise HSV-1 capsids in infected cells^34–36^. However, TEM and cryoSXT have different strategies for introducing contrast in imaging. In TEM, signal is produced by adding a contrast agent, whereas cryoSXT is label-free and contrast is generated via the differential local density of carbon and oxygen in the material. Although cryoSXT has been used previously to image virus particles in infected cells, it was unclear whether individual ‘naked’ HSV-1 capsids, which are approximately 125 nm in diameter^32, 37–40^, would be large enough and offer sufficient contrast to be observed with this imaging method. To establish a baseline, we grew uninfected HFF-hTERT cells on perforated carbon electron microscopy (EM) grids and plunge cryocooled them for imaging by cryoSXT. Unlike a glass lens that focuses light by refraction, a zone plate was used to focus the X-rays by diffraction: a zone plate is a diffraction grating composed of a series of concentric rings in which alternating rings are transparent to X-rays and the resolution is determined by the diameter of the outermost ring^41^. An objective zone plate with 25 nm outer zone was used for our experiments here, affording image resolution of up to 30 nm. To produce a 3D imaging volume (tomogram), a series of X-ray projection images (tilt series) were collected from a single 9.46×9.46 µm field of view in the cell, with each projection collected following rotation of the specimen around an axis normal to the incident X-ray beam. For each tomogram the projections spanned up to 120° of rotation with increments of 0.2° or 0.5° per image. To correct for small inaccuracies in the tilting of the microscope stage during imaging, the projections in the series were aligned together in the program IMOD^42^ using gold fiducials or lipid droplets as landmarks for registration. We collected 19 tilt series that were processed into 3D tomograms. We found that uninfected cell nuclei lacked distinctive internal features other than a difference in average intensity when compared with the cytosol (Fig 1A).

**Fig. 1.**
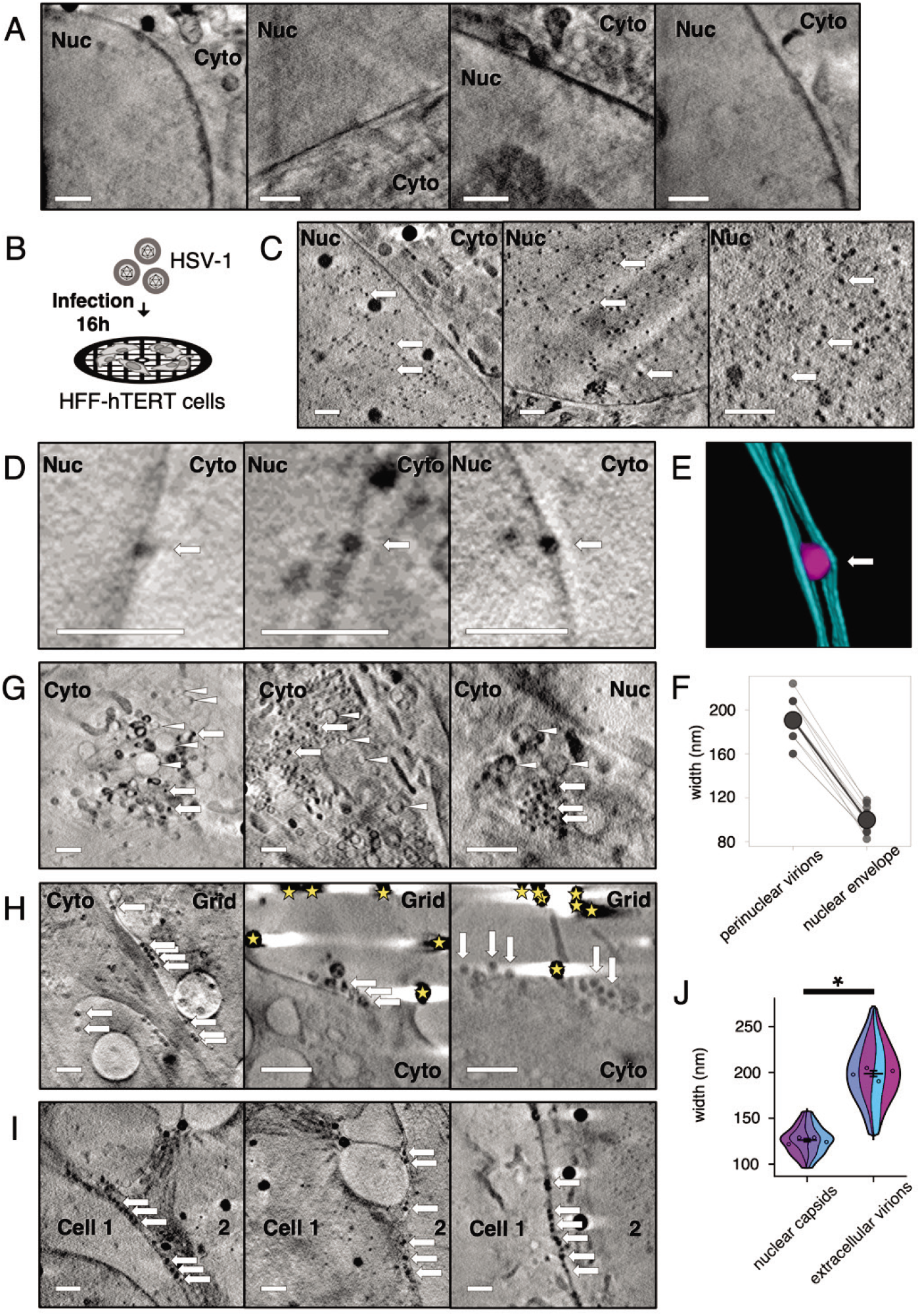
Soft X-ray tomography imaging at cryogenic temperatures of HSV-1-infected HFF-hTERT cells identifies virus particles. HFF-hTERT cells were grown on EM grids, infected (MOI 2) with HSV-1 or mock-infected, and plunge cryocooled 16 hpi. All tomograms were reconstructed from X-ray projections collected using 25 nm (**A**) or 40 nm (**C, D, G–I**) zone plate objectives; scale bars = 1 µm. (**A**) The nucleus (Nuc) has a largely uniform X-ray absorbance in uninfected HFF-hTERT cells. Cyto, cytoplasm. (**B**) Schematic of infection workflow. (**C**) In HSV-1 infected cells many dark puncta are evident in the nucleus, consistent with these puncta being newly assembled HSV-1 capsids. (**D**) Dark puncta were also observed within the perinuclear space of the nuclear envelope, consistent with these being HSV-1 capsids undergoing primary envelopment/de-envelopment to leave the nuclear space. (**E**) Segmentation of a perinuclear viral particle (magenta) and the two membranes of the nuclear envelope (cyan). The perinuclear viral particle expands the nuclear envelope. (**F**) The width of perinuclear viral particles plus associated membranes is 190.5 ± 6.01 nm SEM (N=11; 20.8 nm SD), which is greater than the width of the nuclear membrane elsewhere (99.8 ± 3.57 nm SEM; N=11; 11.9 nm SD). (**G**) HSV-1 capsids (arrows) were also observed in the cytoplasm alongside vesicles (arrowheads). (**H**) Multiple particles are observed along the surface of infected cells, consistent with these being assembled HSV-1 virions that have exited the infected cell. Gold fiducials are indicated with stars. (**I**) HSV-1 virions are also observed at the junctions between cells. (**J**) The width of the nuclear capsids is 125.8 ± 1.70 nm SEM (n=80 from 4 tomograms), consistent with these being HSV-1 capsids (∼125 nm^37, 38^). The width of the extracellular virions is 198.6 ± 3.48 nm SEM (n=80 from 4 tomograms), consistent with these being fully-enveloped HSV-1 virions (∼200 nm^51^). Due to unequal variance, a Mann-Whitney *U* test was performed to determine a significant difference in the width of nuclear capsids and extracellular virions (W=126, *p*-value<2.2×10^-16^). Error bars show mean ± SEM.

Given that the nucleus is the site of capsid assembly, we sought to determine whether an abundance of capsids could be detected in infected cells. To this end, HFF-hTERT cells were cultured on perforated-carbon EM grids, infected with HSV-1 at a multiplicity of infection (MOI) of 2 and plunge cryocooled at 16 hours post-infection (hpi). Infected cells were imaged via cryoSXT using a 40 nm zone plate objective, illuminating a 15.14×15.14 µm field of view, using the image acquisition and analysis workflow detailed above (Fig. 1B). These samples were prepared and cryopreserved on three separate occasions and 98 tomograms were collected in total. Numerous dark puncta were observed in the nucleus of infected cells (Fig. 1C). We interpreted these puncta to be HSV-1 capsids because capsids are rich in carbon and phosphorous, being proteinaceous shells surrounding tightly packed DNA genomes, and these elements exhibit strong absorption at the 500 eV X-ray energy used here for imaging^29^.

During virus assembly, capsids enter the perinuclear space by budding at the inner nuclear membrane (primary envelopment), forming a membrane-wrapped perinuclear virus particle that rapidly fuses with the outer nuclear membrane *en route* to the cytoplasm^9^. These enveloped virions in the perinuclear space are infrequently observed by EM^5, 43–46^ because they are short-lived and the thin sectioning required for imaging using electrons decreases the probability that such structures will be present within the cellular volume being examined. The penetrating power of soft X-rays in unstained cryopreserved samples (> 10 μm in depth) removes the requirement for sectioning, allowing the entire depth of the cell to be imaged for any given field of view. This increases the likelihood of observing short-lived structures such as primary enveloped virus particles. Dark puncta within the nuclear envelope that are likely to be perinuclear viral particles were found 11 times in 98 tomograms (Fig. 1D). The perinuclear viral particles appear to expand the perinuclear space and the nuclear envelope, as shown in a segmented image (Fig. 1E). The width of the nuclear envelope at putative sites of primary envelopment (190.5 ± 6.01 nm SEM; N=11) is significantly greater than the width of the nuclear envelope in other places on the same tomograms (99.8 ± 3.57 nm SEM; N=11; paired t-test p-value=1.93×10^-9^) (Fig. 1F). This demonstrates that substantial deformation of the nuclear envelope must occur to accommodate the presence of perinuclear virus particles.

Dark puncta representing viral capsids were also observed in the cytoplasm in close proximity to vesicles, highlighting potential sites of secondary envelopment (Fig. 1G). After secretion, HSV-1 particles commonly remain bound to the cell surface, a property that may be exacerbated by the antiviral restriction factor tetherin^47, 48^. In addition, we expected to see HSV-1 particles between cells because virions are targeted to cell junctions to promote cell-cell spread^49^. Linear arrays of dark puncta were observed on the cell surface and between cells (Fig. 1H **&** I) and likely represent released virus particles (extracellular virions). Virus particles increase in size during the assembly process as they accumulate their tegument and become enveloped in the cytoplasm before they are released from the cell. We measured the width of nuclear capsids and extracellular virions from 8 tomograms to determine if they could be distinguished based on their size (Fig. 1J). Nuclear capsids had a width of 125.8 ± 1.70 nm SEM (n=80 from 4 tomograms; range 96–160 nm; SD 15.22 nm), which is consistent with high-resolution structural analysis of purified capsids^38^ (∼125 nm) and of capsids inside infected-cell nuclei^50^. Extracellular virions were larger with a width of 198.6 ± 3.48 nm SEM (n=80 from 4 tomograms; range 128–272 nm; SD 31.15 nm), consistent with previous reports (∼175-200 nm^37, 51^). These differences were found to be significant with a Mann-Whitney *U* test for unequal variance (W=126, *p*-value<2.2×10^-16^).

### Fluorescently tagged ICP0 and gC can be used to monitor the progression of HSV-1 infection in HFF-hTERT and U2OS cells

Recent microscopy and single-cell transcriptomics studies have revealed that, even in a monolayer of cultured cells synchronously infected with HSV-1, individual cells progress through the infection cycle at different rates and the remodelling of cellular compartments varies depending on the stage of infection^16, 52^. To control for this, a recombinant strain of HSV-1 termed the timestamp virus has been developed to allow identification of the stage of infection based on the abundance and subcellular localization of the fluorescently tagged early and late viral proteins ICP0 and gC, respectively^16^. Fluorescence microscopy of HFF-hTERT cells infected with this timestamp virus allowed characterization of the changes to cellular compartments that accompany progressing HSV-1 infection and categorization of cells into 4 stages of infection. Having confirmed that virus particles could be observed in infected cells using cryoSXT, we sought to obtain higher-resolution temporal information on the morphological changes that occur over the course of HSV-1 infection by using the timestamp virus. Preliminary experiments performed using infected HFF-hTERT cells were unsuccessful as they proved sensitive to prolonged exposure to the soft X-ray beam when collecting data with a 25 nm zone plate, the objective available at the time on the microscope at the synchrotron beamline used for these experiments, leading to localized sample heating and low-quality tomograms. We thus turned instead to U2OS osteosarcoma cells, which have been used previously for HSV-1 ultrastructural analysis^53, 54^ and have been shown previously to be robust imaging subjects that yield consistently high-quality tomograms when exposed to high doses of soft X-rays^32, 33^.

To compare the temporal profiles of progression of timestamp HSV-1 infection in HFF-hTERT and U2OS cells, we first compared the expression patterns of the fluorescently tagged proteins between the two cell types. Cells were infected at an MOI of 1–3 and samples were fixed at multiple time points following infection before imaging on a widefield fluorescence microscope (Fig. 2). The immediate-early HSV-1 protein ICP0 was used to characterize early stages of infection because it is one of the first viral proteins to be expressed^55^. In both cell lines, eYFP-ICP0 was expressed throughout the course of infection. However, the spatial localization of eYFP-ICP0 differed somewhat between HFF-hTERT and U2OS cells. In HFF-hTERT cells, eYFP-ICP0 was observed in the nucleus in stage 1 whereas it became more concentrated in the cytoplasm with relatively weaker signal in the nucleus in stages 2–4. However, in U2OS cells, eYFP-ICP0 expression displayed a high signal in the nucleus throughout infection while also becoming more concentrated in the cytoplasm as infection progressed, suggesting U2OS cells retain more eYFP-ICP0 in the nucleus at the later stages of infection than is observed for HFF-hTERT cells (Fig. 2A). This may reflect differences in cellular interactions for ICP0 in U2OS cells, which is consistent with previous observations demonstrating that replication deficits demonstrated by ICP0-null strains of HSV-1 in human fibroblasts are effectively complemented in U2OS cells^56^. The continued high signal levels of eYFP-ICP0 in the nucleus complicated the distinction between the early stages (stages 1+2) of infection in U2OS cells.

**Fig. 2.**
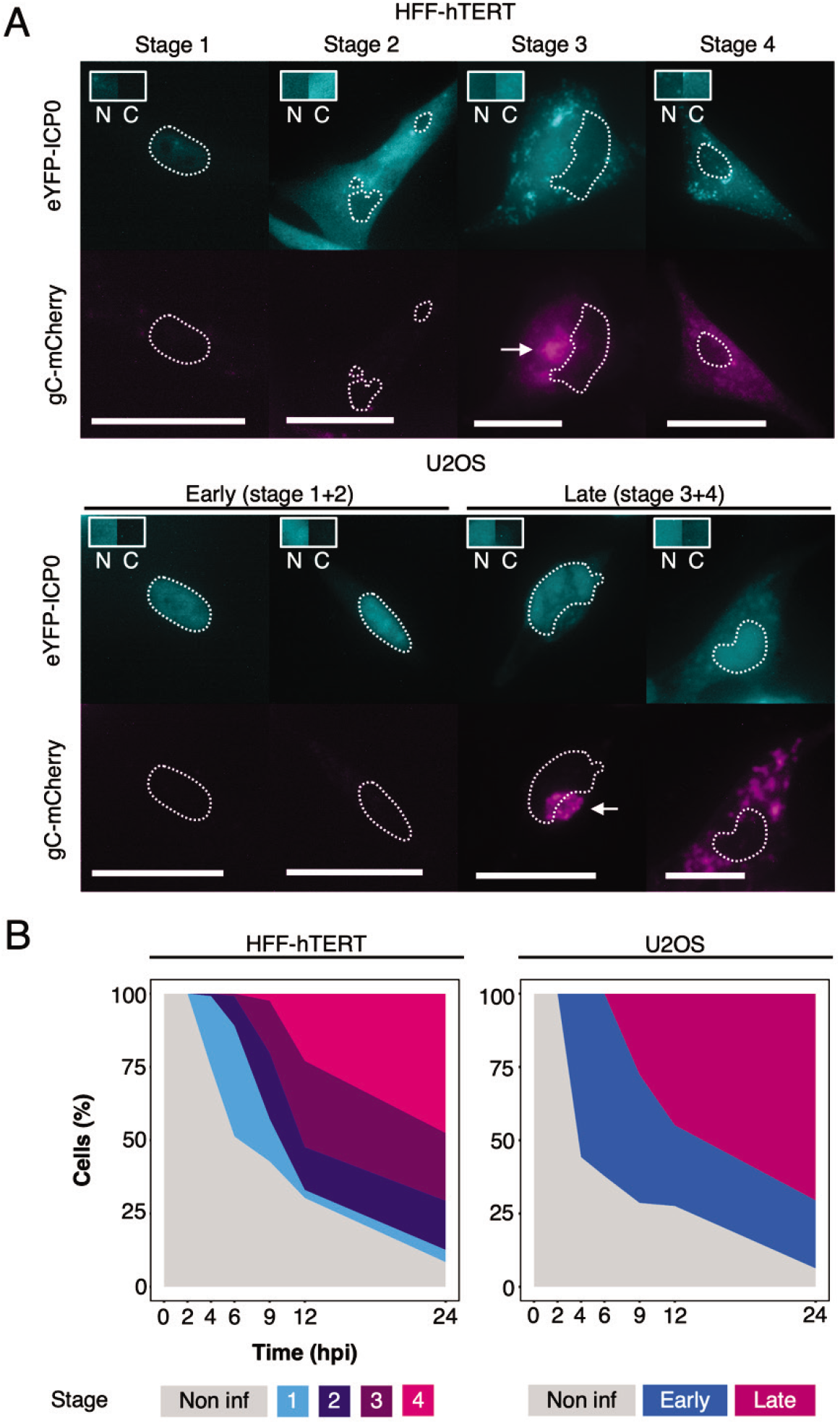
Temporal analysis of HSV-1 infection using the dual-fluorescent timestamp virus. (**A**) Room temperature widefield fluorescence imaging of timestamp HSV-1 infected HFF-hTERT and U2OS cells was used to delineate between stages of infection based on the expression and localization of the early protein eYFP-ICP0 and the late protein gC-mCherry^16^. The spatiotemporal expression of these fusion proteins was similar in HFF-hTERT and U2OS cells, except for increased retention of eYFP-ICP0 in the nucleus of U2OS cells during all stages. Outlines show the nuclei and arrows indicate juxtanuclear compartments rich in gC-mCherry. Scale bars = 50 µm. Boxes show a sample of the eYFP-ICP0 intensity from the nucleus (N) and cytoplasm (C). (**B**) The proportion of infected cells in each stage was determined using widefield imaging at 2, 4, 6, 9, 12, and 24 hpi following infection (MOI 3) of HFF-hTERT and U2OS cells with timestamp HSV-1.

The spatial expression of gC-mCherry was broadly similar between HFF-hTERT and U2OS cells. gC is a viral glycoprotein expressed at late stages of virus replication^57^ that is incorporated into nascent virus particles at sites of virus envelopment^58^. In HFF-hTERT cells, gC-mCherry was enriched at a juxtanuclear site in stage 3 but became fragmented and dispersed throughout the cytoplasm and at the plasma membrane by stage 4 (Fig. 2A). A similar spatial expression was observed for late stage U2OS cells, with redistribution of gC from juxtanuclear sites to the periphery likely representing progressively later stages of infection. However, there existed a continuum of gC distribution between juxtanuclear and dispersed in late stages of infection in U2OS cells. This, combined with the difficulties in differentiating between early infection stages due to nuclear retention of eYFP-ICP0, led us to group U2OS infection stages into two broader categories (“early” and “late”). The designation of early or late-stage infection was determined by the absence or presence of gC-mCherry signal in eYFP-ICP0 positive cells, respectively.

Next, we probed whether progression through the replication cycle follows the same timecourse in HFF-hTERT and U2OS cells. Both cell types were inoculated with timestamp virus (MOI 3) for one hour, at which time unabsorbed viruses were inactivated with a citric acid wash, and cells were fixed at various time points over the course of 24 hrs before imaging (Fig. 2B). We observed that the infection proceeds at a similar pace in both cells types, with a similar proportion of cells in equivalent stages of infection (1+2/early and 3+4/late) at each time point.

### CryoSXT following infection with ‘timestamp’ HSV-1 allows temporal correlation of ultrastructural changes during infection

To characterize the changes in morphology of cellular compartments that accompany different stages of virus infection, U2OS cells were grown on perforated-carbon EM grids before being infected (or mock infected) with timestamp HSV-1 and cryogenically preserved by plunge cryocooling in liquid nitrogen-cooled liquid ethane (Fig. 3A). Vitrified samples were analysed by cryo-widefield microscopy to classify the stage of infection and then imaged using cryoSXT to correlate the stage of virus infection in a specific cell with observed morphological changes (Fig. 3B). Cells were infected at MOI 1 and the samples were plunge cryocooled at 9 hpi in an attempt to evenly sample the different stages of infection (Fig. 2B). In total, 139 tomograms were reconstructed; 76 from uninfected cells alongside 22 and 41 from cells at early or late stages of infection, respectively, across three independent replicates (Table 1). Manual inspection of the resultant tomograms revealed that the 25 nm zone plate allows detection of higher resolution features than is possible with the 40 nm zone plate, such as the lumen of the endoplasmic reticulum, cytoskeletal filaments, and small membrane structures (Supp. Fig. 1A-E). The observed width of nuclear capsids in U2OS cells imaged using the 25 nm zone plate (Supp. Fig. 1F) is similar to those observed in infected HFF-hTERT cells imaged using the 40 nm zone plate (Fig. 1J). The tomograms collected from U2OS cells using the 25 nm zone plate were thus deemed suitable for identifying changes to cellular compartments that occur during HSV-1 infection.

**Fig. 3.**
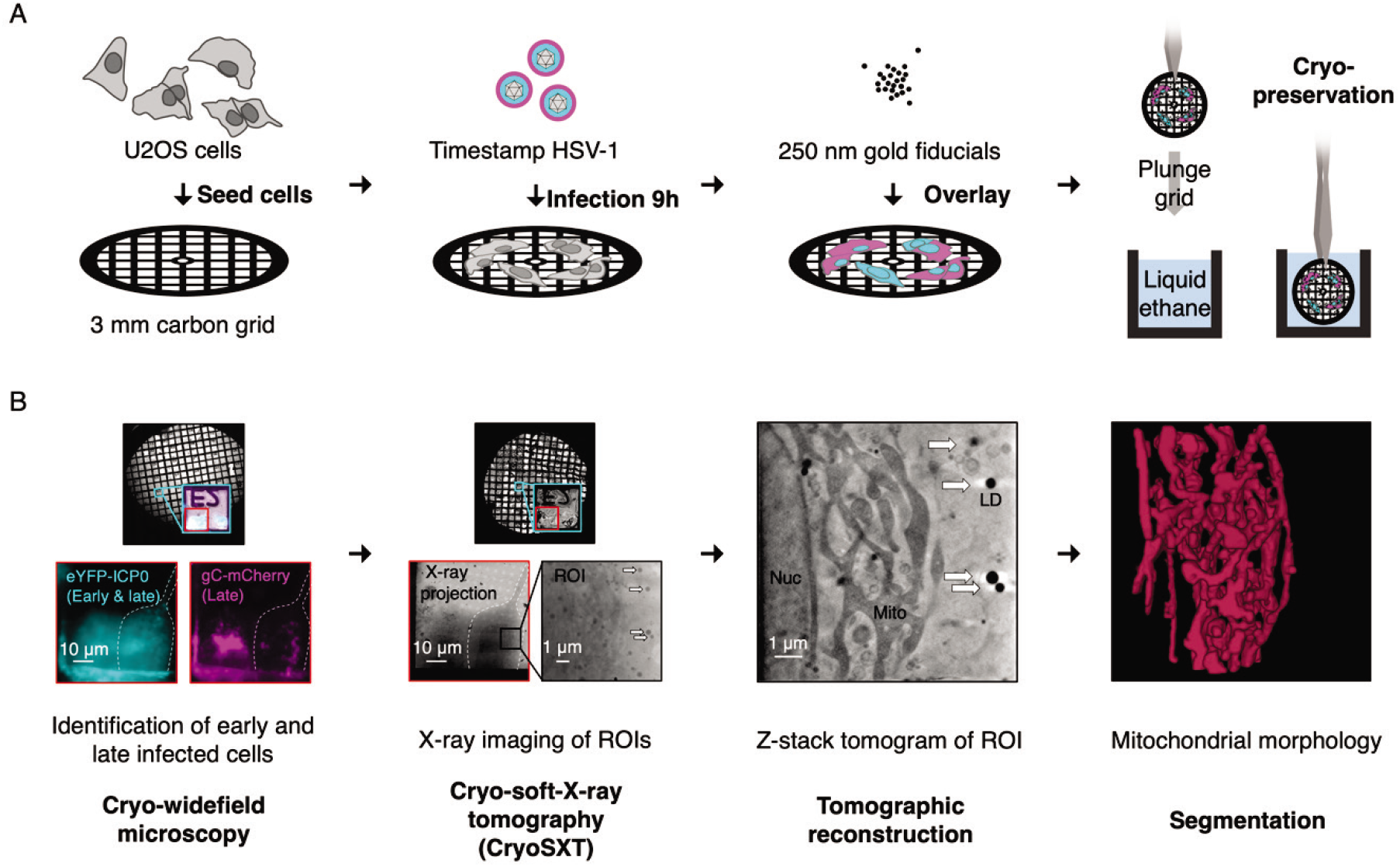
Workflow for multi-modal imaging of HSV-1 infected cells. (**A**) Preparation of infected cells samples for multimodal imaging. U2OS cells are cultured on perforated EM grids and infected with recombinant ‘timestamp’ HSV-1, expressing fluorescently tagged proteins eYFP-ICP0 and gC-mCherry that allow identification of the stage of infection for each cell under investigation. At 9 hpi, gold fiducials are overlayed onto the sample to facilitate image registration and grids are cryopreserved in a near-native state by plunge cryocooling in liquid ethane. (**B**) Multi-modal imaging of infected U2OS cells. A widefield microscope with a cryo stage is used to locate the grid positions of infected cells. The stage of infection for each cell is determined based on the expression of eYFP-ICP0 and gC-mCherry (as shown in Fig. 2). These grids are then loaded into the cryo-soft-X-ray microscope at Diamond Light Source beamline B24 and are illuminated with soft X-rays at the marked grid positions. X-ray projections of regions of interest (ROIs) are collected at multiple angles and aligned using the gold fiducials and intracellular features, such as lipid droplets (LDs), with the program IMOD^42^. Tomograms are reconstructed from these projections using IMOD to compare intracellular morphology between uninfected cells and those at early- or late-stages of infection. Segmentation with tools like *Contour*^60^ facilitates quantitation and visualization in three dimensions of the observed cellular structures.

**Table 1.** Collection of cryoSXT data to analyse changes in cellular morphology accompanying infection. CryoSXT data was collected using a 25 nm zone plate from multiple uninfected cells or cells at early and late stages of infection across three independent replicates. Tiled X-ray projections (‘X-ray mosaics’) with a 66.2×66.2 µm field of view were collected at multiple areas on the sample grid to identify cells of interest. Tilt series were collected at perinuclear or peripheral regions of the cytoplasm within these cells and were processed to generate tomograms.

| Replicate | Stage of infection | X-ray mosaics | Cells in mosaics | Cells imaged by tomography | Tomograms |
| --- | --- | --- | --- | --- | --- |
| 1 | Uninfected | 19 | 30 | 18 | 29 |
|  | Early | 4 | 4 | 2 | 4 |
|  | Late | 8 | 13 | 10 | 14 |
| 2 | Uninfected | 10 | 20 | 14 | 20 |
|  | Early | 9 | 13 | 11 | 13 |
|  | Late | 10 | 10 | 8 | 12 |
| 3 | Uninfected | 8 | 27 | 26 | 27 |
|  | Early | 6 | 7 | 5 | 5 |
|  | Late | 8 | 13 | 13 | 15 |

We observed that HSV-1 infection does not dramatically affect the morphology of the nucleus or integrity of the nuclear envelope, despite the continuous budding and fusion of capsids that occur at the inner and outer nuclear membranes, respectively, during infection. We occasionally observed bulging of the nuclear envelope into the cytoplasm without separation of the inner and outer nuclear membranes (Supp. Fig. 1G). This bulging could be seen in both uninfected and infected cells. It was distinct from the separation of the inner and outer nuclear membranes caused by the expansion of the perinuclear space, which we observed in the presence of perinuclear viral particles (Fig. 1D–F). It also differed from the expansion of the perinuclear space observed previously in uninfected murine adenocarcinoma cells imaged using cryoSXT^59^. The cryopreservation protocol used for preparing our cryoSXT samples does not result in dehydration artefacts that can alter the apparent morphology of the nuclear membrane in TEM samples^59^, suggesting that these bulges are not artefacts of the sample preparation, but the biological relevance of this observation remains unclear.

Striking changes were observed in the size and dispersal of vesicles during HSV-1 infection (Fig. 4 A). HSV-1 capsids are thought to interact with several types of vesicles in the cytoplasm, including *trans*-Golgi network vesicles and endosomes, both of which have been implicated in secondary envelopment^13^. Infected cells had a greater number of vesicles in juxtanuclear regions when compared with uninfected cells (Fig. 4A **and Supp. Video 2**). To determine if there was a difference in the size of vesicles between uninfected cells and those at early- or late-stages of infection we developed *Contour*, a program to segment and quantitate cellular features in 3D volumes^60^. The widest point of each vesicle in three dimensions from 4 tomograms for each condition was measured (Fig. 4B). The mean vesicle width was higher for uninfected cells (802.23 ± 348.47 nm SD, N=96) than for early-stage (688.66 ± 271.76 nm SD, N=184) and late-stage (631.85 ± 270.60 nm SD, N=184) infected cells. The mean vesicle widths for each tomogram were compared using a one-way ANOVA and Tukey test and the vesicle widths of uninfected cells were found to be significantly different from early-stage (p=0.04) and late-stage (p=0.01) infected cells. The vesicle width did not differ significantly between early-stage and late-stage infected cells (p=0.62). Segmented vesicles (Fig. 4A) were open-ended because the contrast of membranes in the sample differs based on their orientation with respect to incident X-ray beam. This arises because the sample can only be rotated by 120° during cryoSXT tilt series image acquisition, rather than 180° as would be required for isotropic data collection, due to geometric constraints between microscope and sample holder components. The edges of vesicles that lie parallel to the incident X-ray beam (i.e. the ‘sides’ of the vesicle with respect to the XY projection plane) produce high contrast, since the X-rays pass tangentially through the carbon-rich membrane of the vesicle and thus traverse a large volume of material that strongly absorbs X-rays. The ‘front’ and ‘back’ edges of the vesicle with respect to the XY projection plane yield less contrast because the X-rays pass radially through these membranes, traversing a shorter path through this carbon-rich X-ray absorbing material. The lower contrast for the front and back edges of vesicles prevented their reliable segmentation, yielding gaps in the resultant volumes.

**Fig. 4.**
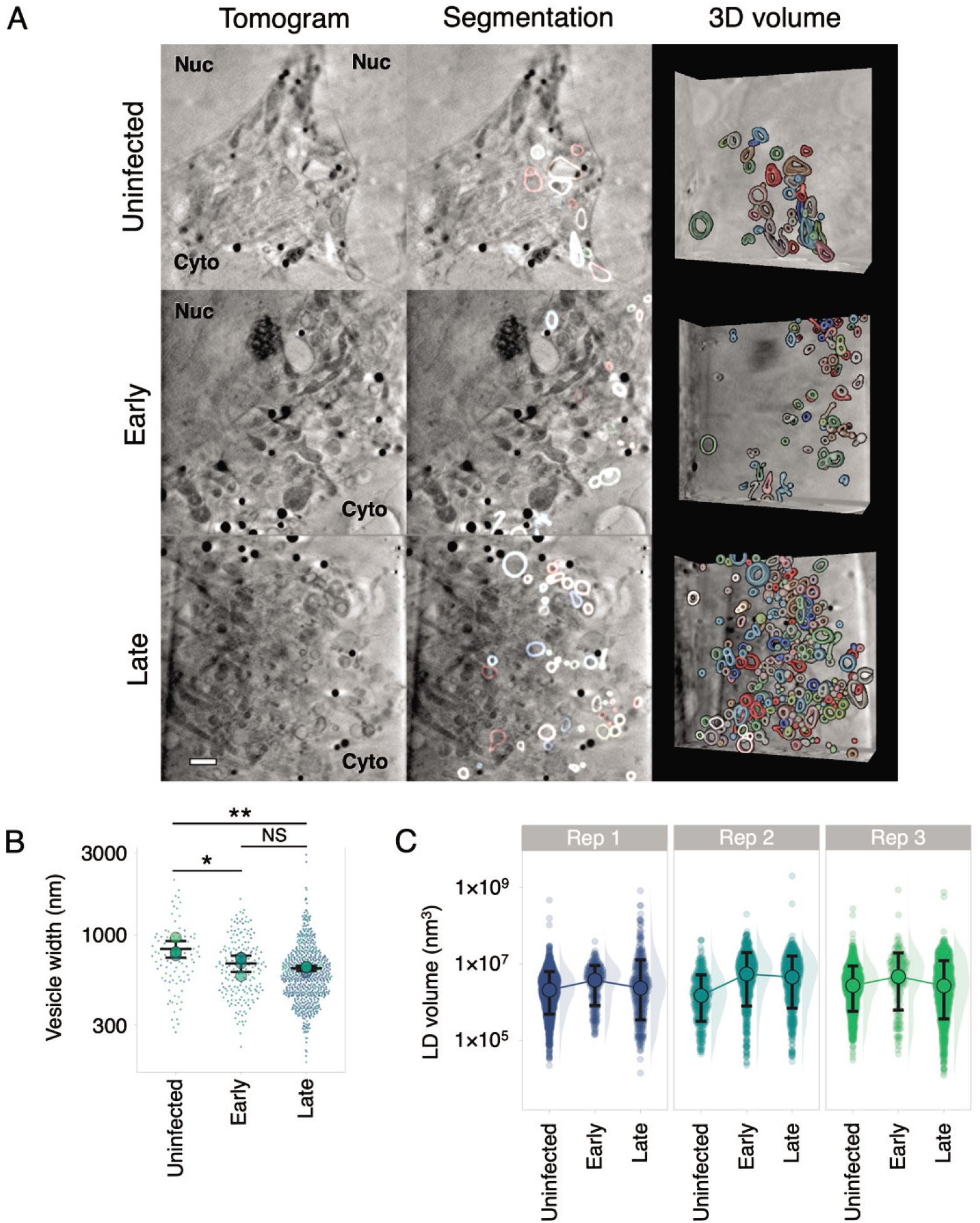
Remodelling of cytoplasmic vesicles during HSV-1 infection. CryoSXT tomograms were recorded from uninfected cells, or cells at an early or late stage of infection with timestamp HSV-1, as determined via wide field fluorescence cryo-microscopy. Data are representative of three independent experiments. Scale bar = 1 µm. (**A**) A higher concentration of vesicles is observed at the juxtanuclear compartment in cells at early- or late-stages of infection compared with uninfected cells. (**B**) The maximum width of each vesicle in three-dimensions was measured in *Contour*^60^. Width was measured instead of volume because the segmented vesicles were open-ended owing to reduced contrast in the tomograms of membranes normal to the incident X-ray beam. Vesicles with a spherical, ellipsoidal, or dumbbell shape were included in the analysis but vesicles with a shape that didn’t fall into these categories were excluded. Intra-luminal vesicles and vesicles that were not individually resolved by the segmentation were also excluded from the analysis. Significance of differences was assessed with a one-way ANOVA and Tukey tests for the combinations: uninfected-early (p=0.04), uninfected-late (p=0.01), and early-late (p=0.62). Big circles show the mean vesicle width per tomogram (4 tomograms per condition). Error bars show overall mean ± SD. (**C**) Lipid droplets were segmented and measured using *Contour*^60^ and their distributions were plotted on a logarithmic scale. Median volumes ± SD (hollow circles plus error bars) are shown for each group because median values are less affected than mean values by non-normal distributions. The median volume was highest in cells at early stages of infection in all three replicates. A linear plot of the distributions and significance tests for the lipid droplet volumes are shown in Supp. Fig. 2.

Lipid droplets are carbon-dense organelles that produce high contrast in cryoSXT and were clearly visible as dark solid spheres in the tomograms. These lipid droplets could be readily segmented using *Contour*^60^, allowing measurement of their volumes. We observed an increase in the volume of these droplets in cells during the early stage of infection when compared with uninfected cells or to cells at the late stage of infection (Fig. 4C). A small number of extremely large lipid droplets (>5×10^7^ nm^3^) were observed in tomograms from each of the three replicate infections, but the presence of these large droplets was not correlated with progression through the infection (Supp. Fig. 2A-B). The distribution of lipid droplet sizes was non-Gaussian (positively skewed; Supp. Fig. 2C) and a non-parametric Mann-Whitney *U* test confirmed that, in all three replicate experiments, the lipid droplets were significantly larger in cells at early stages of infection than in uninfected cells (Supp. Fig. 2D). There is not a consistent difference in the size of lipid droplets between uninfected cells and those at late stages of infection, suggesting that lipid droplets undergo a transient change in size during infection.

Mitochondria were the most phenotypically diverse organelles monitored in this study. In most cases, they were thin and possessed a dark matrix (Fig. 5A). However, occasionally there were cells that contained swollen mitochondria with a lighter matrix with highly contrasting cristae (Supp. Fig. 3A), similar to observations of mitochondria made by EM^61–63^ and cryoSXT^59^. This swollen morphology can be associated with release of cytochrome *c* from porous mitochondria during apoptosis^61^. Swollen mitochondria were observed in each of the three independent sets of cell growth, infection and plunge cryocooling experiments performed, but these swollen mitochondria were most prevalent in the uninfected cells of replicate 3 (Supp. Fig. 3B). In uninfected cells, non-swollen mitochondria were heterogeneous in shape, with numerous being small and spherical or long and curved in the same cell (Fig. 5A). We observed branching in some elongated mitochondria. However, mitochondria appeared less heterogenous in shape in infected cells, and were consistently more elongated and branched (Fig. 5B-D, Supp. Fig. 3C, and **Supp. Video 3**), in line with previous observations made using super-resolution fluorescence microscopy of HFF-hTERT cells infected with the timestamp virus^16^. The number of points where mitochondria branch into two or more arms (branching nodes) was significantly increased (p<0.05) in cells at late stages of infection (20.5 ± 5.45 nodes SD; n = 15) compared with uninfected cells (7.0 ± 4.02 nodes SD; n = 15) according to a one-way ANOVA and Tukey tests performed on each replicate (Fig. 5D). In some cases, the mitochondria fused into a single, branched network (Fig. 5B and **Supp. Video 3**), providing a dramatic demonstration of the increase in mitochondrial branching and decrease in number of distinct mitochondrial networks that accompanies HSV-1 infection. It was also observed that the number of distinct mitochondria decreased in infected cells, although ambiguity regarding the connectivity of mitochondrial networks that extend beyond the tomogram field-of-view prevented precise quantitation of this effect. Confocal microscopy qualitatively confirmed the observations made with cryoSXT that mitochondria appear more elongated in infected cells and that mitochondrial morphology in peripheral areas of the cell was more heterogenous in uninfected cells than in infected cells (Supp. Fig. 4). However, the limited resolution of confocal imaging made it difficult to differentiate between highly-branched and separate-but-overlapping mitochondria, particularly at crowded juxtanuclear locations.

**Fig. 5.**
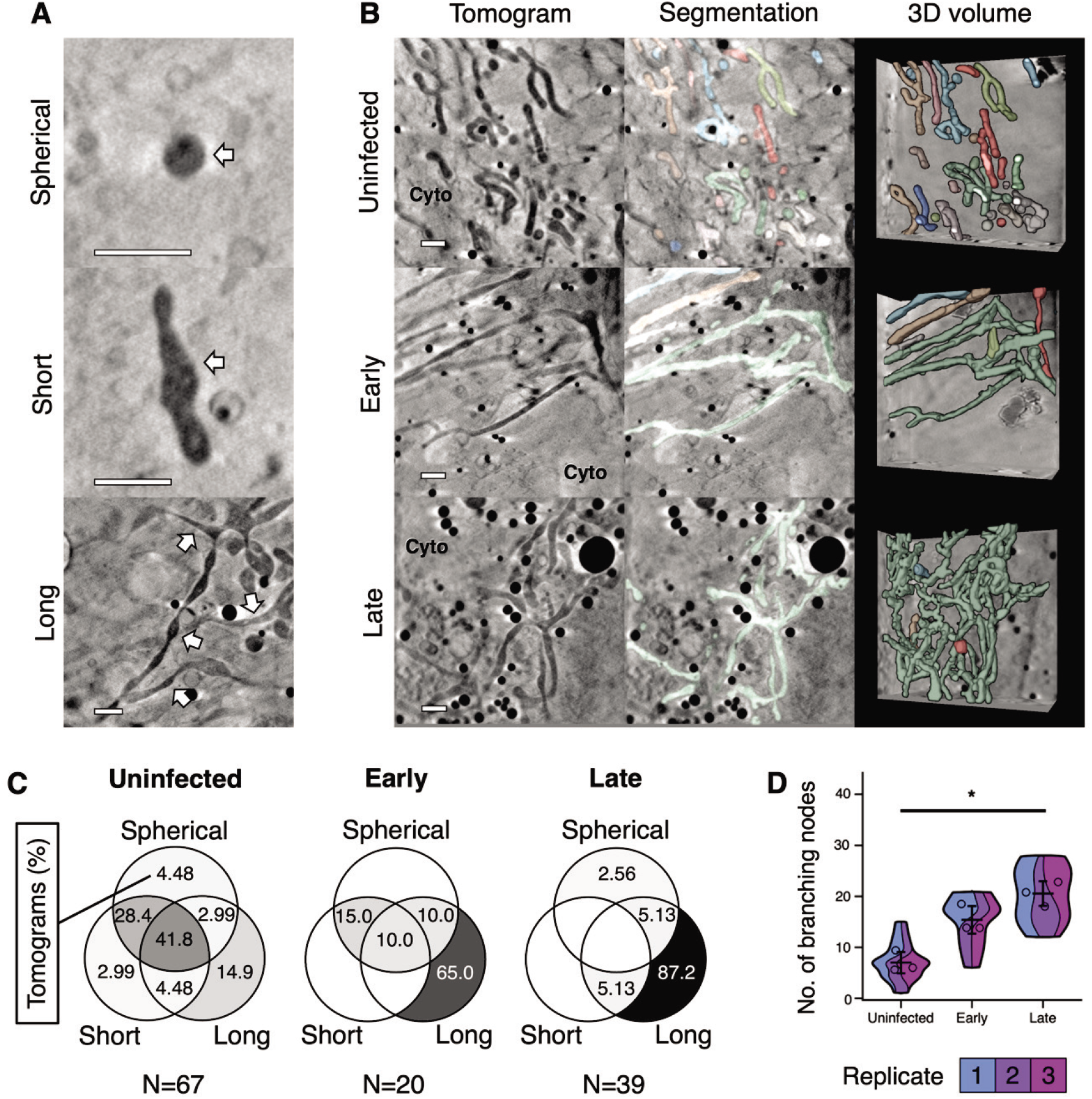
Remodelling of mitochondria during HSV-1 infection. Morphological changes to mitochondria were assessed from cryoSXT tomograms collected from uninfected cells and cells at early- or late-stages of infection with timestamp HSV-1. Data are representative of three independent experiments. Scale bars = 1 µm. (**A**) Examples of spherical, short, and long mitochondria are indicated with white arrows. (**B**) A shift towards elongated and branched mitochondria was observed during infection. Mitochondria were segmented and differentiated using *Contour*^60^ to highlight the abundance and 3D geometry of individual mitochondria. (**C**) Venn diagrams showing the percentage of tomograms at each stage of infection with Spherical, Short or Long mitochondria, or a combination of these phenotypes. The percentages of tomograms with long mitochondria were greater for cells at early- or late-stages of infection than for uninfected cells. Mitochondrial morphology was more heterogenous in uninfected cells. Combined percentages from all replicates are shown here and Venn diagrams for each replicate are shown in Supp. Fig 2C. (**D**) The numbers of branching nodes were calculated for 45 tomograms across all replicates and significant differences in the number of nodes between uninfected cells and those at late stages of infection were determined for each replicate using ANOVA and Tukey tests (p <0.05). Error bars show mean ± SD.

### Golgi membranes and the microtubule network are disrupted during HSV-1 infection of U2OS cells

HSV-1 infection is known to be accompanied by dispersal of the Golgi complex and fragmentation of the *trans*-Golgi network^16, 64, 65^. However, Golgi-related compartments can be difficult to distinguish from other vesicular compartments by cryoSXT and are infrequently observed^59^. We therefore used SIM super-resolution fluorescence microscopy to monitor the changes in Golgi organisation that accompany HSV-1 infection. Fixed U2OS cells that had been infected (MOI 3) with the timestamp reporter HSV-1 for 6 hours were immunostained with the *cis*-Golgi marker GM130, demonstrating that the GM130^+^ Golgi membranes are clustered with a tubular morphology at early stages of infection (Fig. 6A). In cells at late stages of infection the distribution of GM130 was more punctate and more widely distributed throughout the cell, consistent with fragmentation of the Golgi (Fig. 6B). The gC-mCherry signal was also present at multiple sites throughout the cell, often adjacent to the GM130 signal (Fig. 6B). SIM imaging of infected U2OS cells stained with the *trans*-Golgi network marker TGN46 also demonstrated increasing dispersion of TGN46^+^ membranes at late-versus early-stages of infection (Fig. 7) and again we observed that the TGN46 signal was adjacent to, or partially overlapping with, gC-mCherry signal in cells at late stages of infection (Fig. 7B).

**Fig. 6.**
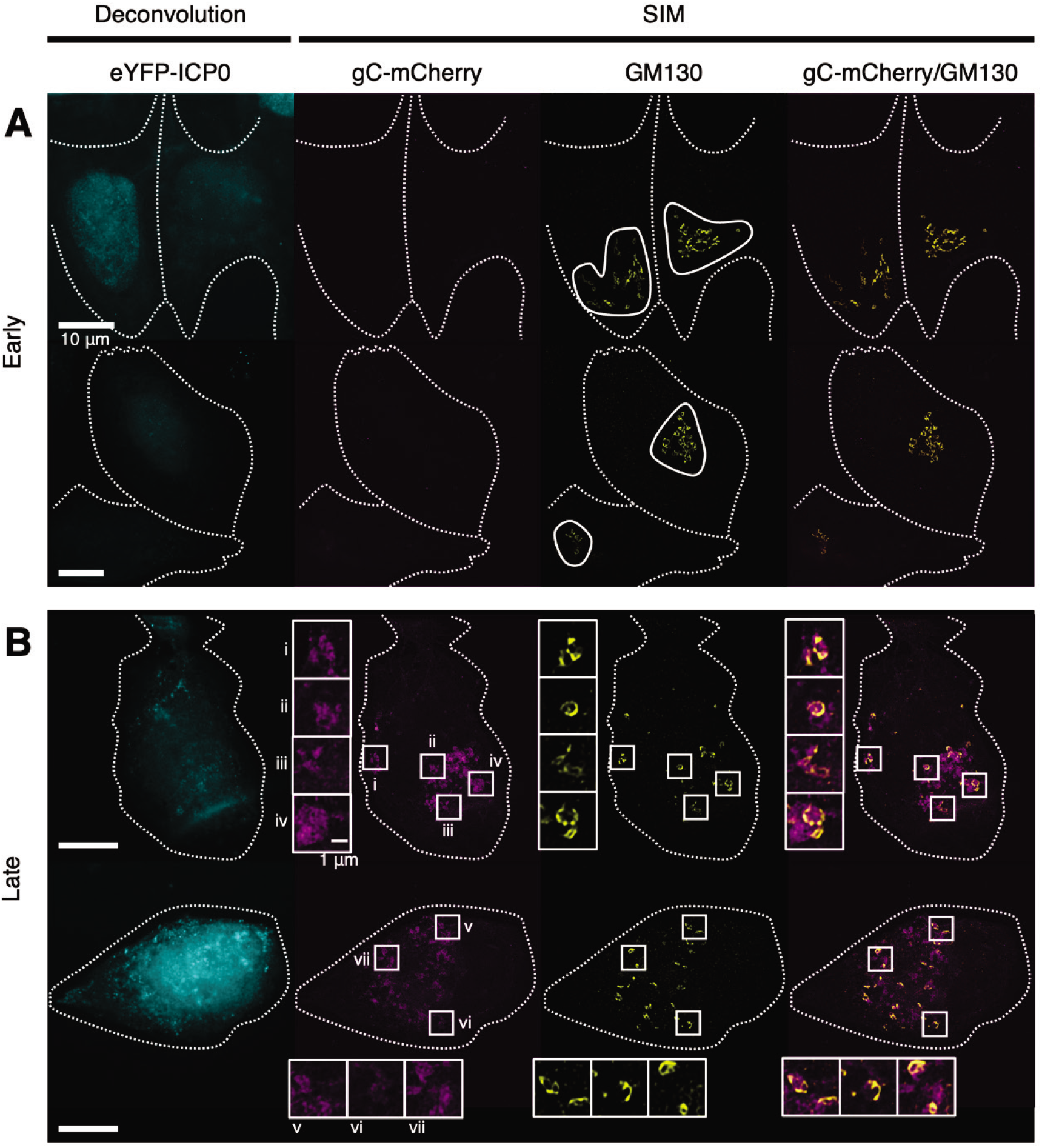
Fragmentation and dispersal of *cis*-Golgi membranes during HSV-1 infection. U2OS cells infected (MOI 3) with timestamp HSV-1 were fixed at 6 hpi and imaged by SIM and deconvolution microscopy. GM130 immunolabelling was used to identify *cis*-Golgi membranes^106^. Dotted outlines denote the cell boundaries. (**A**) Cells at early stages of infection were identified by the presence of eYFP-ICP0 signal using deconvolution microscopy and by the absence of high gC-mCherry signal using SIM. GM130^+^ membranes, which appeared clustered at early stages of infection, are outlined. (**B**) Cells at late stages of infection were identified by the presence of high gC-mCherry signal. GM130^+^ membranes were dispersed and fragmented in these cells. Boxes (i–vii) and corresponding insets showing adjacent localization of GM130 and gC-mCherry.

**Fig. 7.**
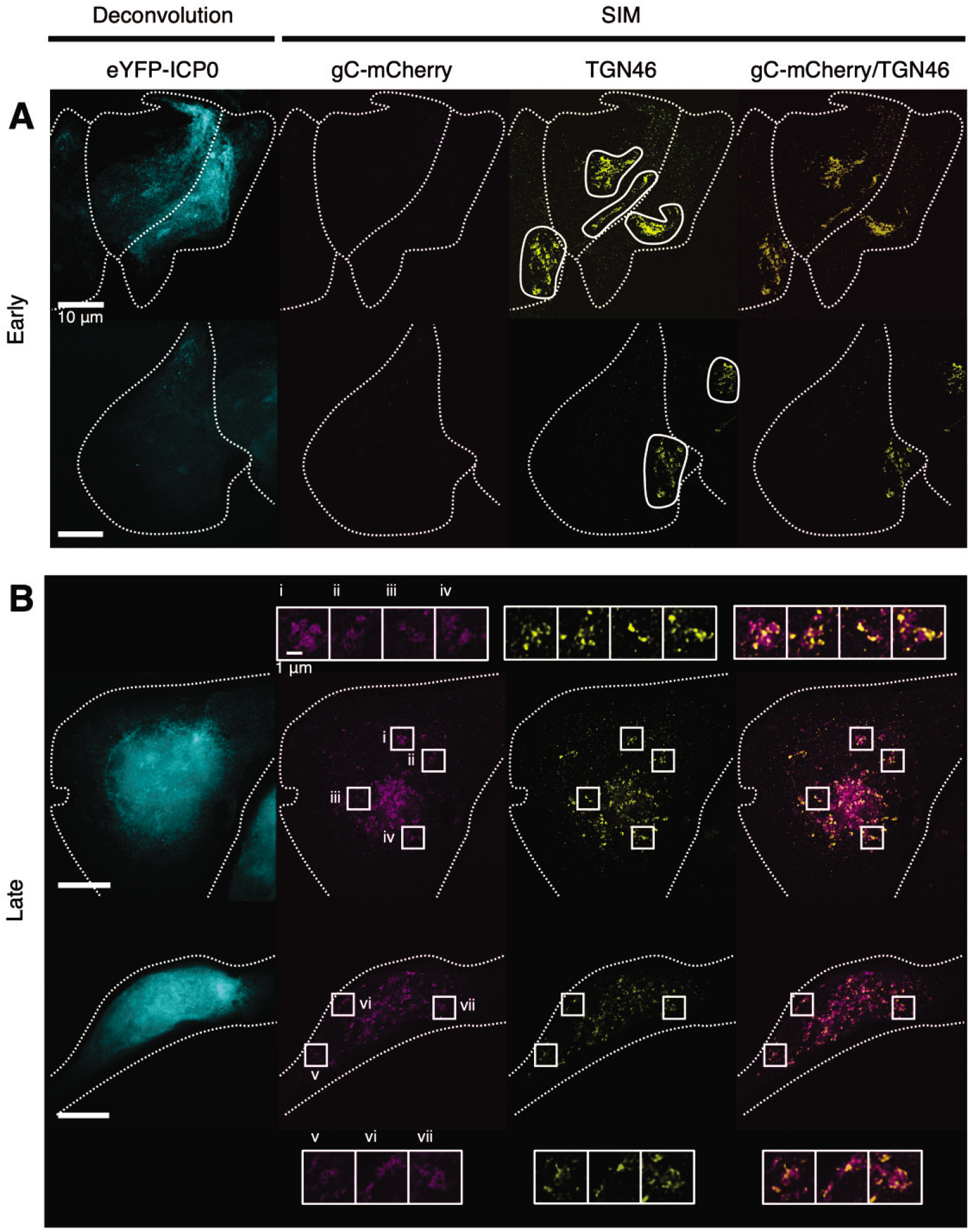
Fragmentation and dispersal of *trans*-Golgi membranes during HSV-1 infection. U2OS cells infected (MOI 3) with timestamp HSV-1 were fixed 6 hpi and imaged by SIM and deconvolution microscopy. TGN46 immunolabelling was used to identify *trans*-Golgi network membranes^107^. Dotted outlines denote the cell boundaries. (**A**) Cells at early stages of infection were identified by the presence of eYFP-ICP0 signal using deconvolution microscopy and by the absence of high gC-mCherry signal using SIM. TGN46^+^ membranes, which appeared both clustered and dispersed at early stages of infection, are outlined. (**B**) Cells at late stages of infection were identified by the presence of high gC-mCherry signal using SIM and TGN46^+^ membranes were widely dispersed in these cells. Boxes (i–vii) and corresponding insets indicate sites of colocalization and adjacent signal between TGN46 and gC-mCherry.

As demonstrated above, HSV-1 infection of U2OS cells changes the morphology of lipid droplets, mitochondria, vesicles and Golgi membranes, all of which interact with the microtubule network^66–69^. While cytoskeletal filaments can occasionally be observed using cryoSXT (Supp. Fig. 1C), microtubules are too thin (25 nm width)^70^ to be reliably detected using this technique. We therefore used confocal fluorescence microscopy to study microtubule morphology in uninfected U2OS cells or cells infected with timestamp HSV-1 (MOI 3) at early and late stages of infection, captured at 6 and 16 hpi, respectively. Microtubules were monitored by immunolabelling β-tubulin. In uninfected cells microtubules formed long filaments that radiated out of a prominent microtubule organising centre (MTOC) (Fig. 8A). By early stages of infection, MTOCs were less pronounced and microtubules no longer had a well-dispersed, radial distribution (Fig. 8B). At late stages of infection, MTOCs could not be detected in most cells, the microtubule network became more compact, and fewer long filaments were observed (Fig. 8C).

**Fig. 8.**
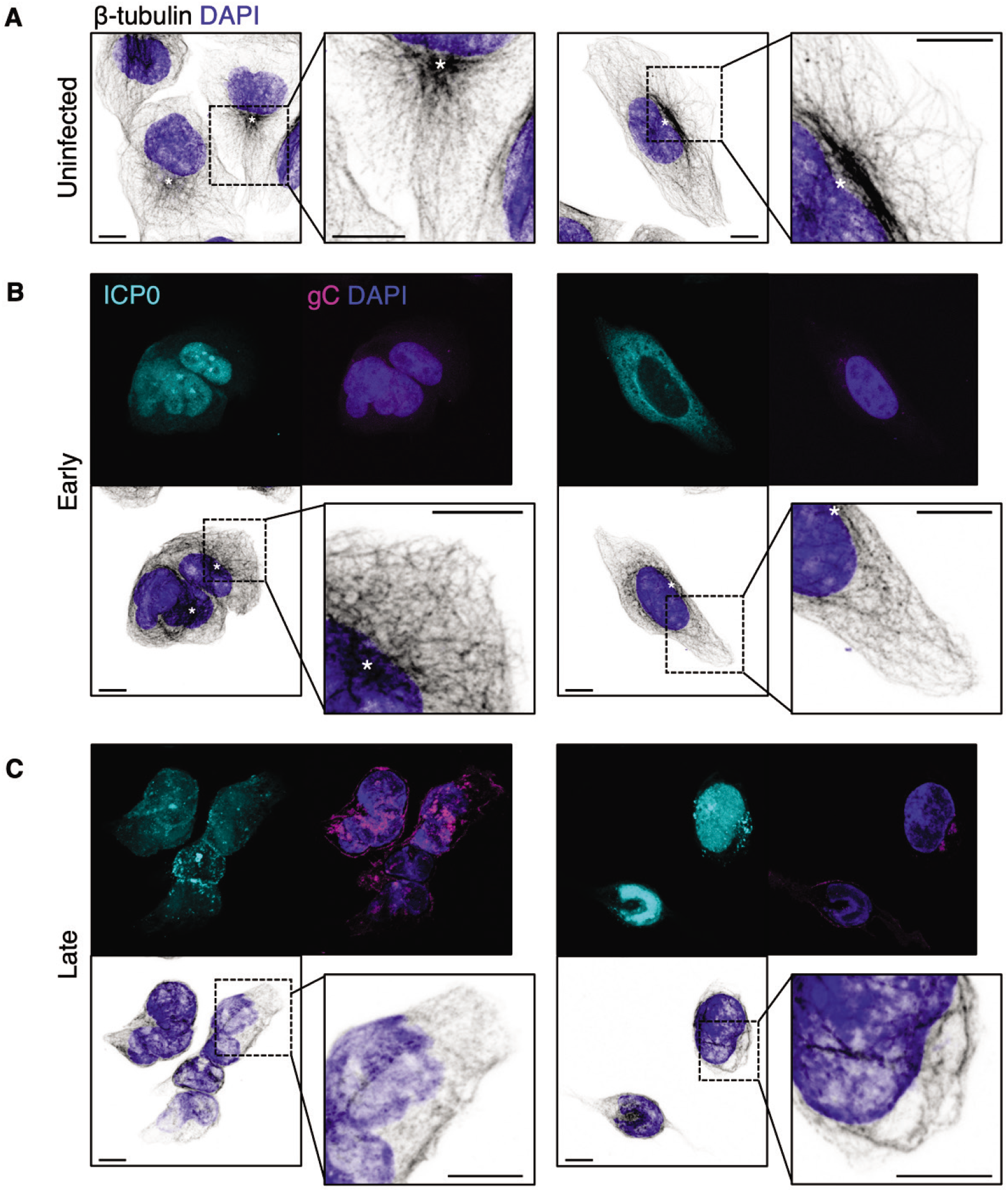
Remodelling of microtubules during HSV-1 infection. U2OS cells infected with timestamp HSV-1 were fixed at indicated times and imaged by confocal microscopy. β-tubulin immunolabelling was used to identify microtubules. Scale bars = 10 μm. Putative microtubule organising centres (MTOCs) are indicated with asterisks (*). (**A**) Uninfected cells exhibited an outspread microtubule network with long filaments, largely radiating from a putative MTOC. (**B**) The microtubule network was closely packed in cells at early stages of infection (6 hpi). (**C**) In cells at late stages of infection (16 hpi), fewer long filaments were observed, the cells lacked noticeable MTOCs, and the network became very closely packed.

## Discussion

In this study, we demonstrated that cryoSXT can be used to monitor the production of nascent HSV-1 particles and observed changes to the architecture of cellular compartments during infection. The sizeable field of view and penetrating power of X-rays facilitate cryoSXT imaging throughout the depth of the cell, allowing rare or transient events to be captured such as the transit of nascent capsids through the nuclear envelope. Furthermore, the lack of requirement for contrast-enhancing agents or chemical fixation allows direct imaging of cellular compartments in a near-native state. We exploited these properties of cryoSXT to compare the morphology of cellular compartments between uninfected and infected U2OS cells, using a recombinant strain of HSV-1 expressing fluorescently tagged early and late viral proteins to identify the infection stage of individual cells within the virus-inoculated samples.

CryoSXT has several advantages as a technique for probing the ultrastructure of cells, plus a number of limitations. CryoSXT imaging is performed on samples that have been vitrified through plunge cryocooling. This rapid and convenient sample preparation technique preserves cellular ultrastructure in a near-native state, avoiding the artefacts that have been associated with chemical fixation, dehydration, and resin-embedding for TEM analysis^29^ and yielding higher label-free contrast than is obtained using cryoET^71^. Furthermore, the penetrating power of X-rays means that samples up to 10 μm thick can be imaged by cryoSXT^29^. As the U2OS cells we investigated had an average depth of approximately 3 μm, each X-ray tomogram contained hundreds of projections through the entire depth of the cell. This contrasts with TEM, cryoEM, and cryoET imaging, which generally require ultra-thin sectioning or focused ion beam (FIB)-milling of samples into ∼0.5–1.0 μm lamella such that the entire depth of an adherent cell like U2OS cannot be collected in one acquisition^72–74^. CryoSXT images a large field of view (9.46×9.46 μm and 15.14×15.14 μm with the 25 nm and 40 nm zone plates, respectively), allowing regions of the nucleus, perinuclear space, and peripheral cytoplasm to be captured together. Lastly, cryoSXT is a relatively high-throughput imaging technique, with each tomographic dataset taking only 5–20 minutes to collect depending on X-ray beam brightness, exposure time, angular rotation per frame and total rotation range used for tomogram acquisition^29^. As we demonstrated in this study, the ability to conveniently prepare samples and collect multiple tomograms expands the number of cells that can be interrogated, allowing robust numerical analysis of the biological specimen under investigation. For example, by using the semi-automated segmentation tool *Contour*^60^ we were able to analyse the volumes of 4845 individual lipid droplets, acquired across 94 tomograms from three biological replicate HSV-1 infections, unambiguously demonstrating a transient increase in lipid droplet volume at the early stage of infection (Fig. 4C, Supp. Fig. 2). We also demonstrate that cryoSXT can be used to perform accurate quantitative analyses of geometric properties of the samples. For example, our measurements of widths of capsids and extracellular virions, determined from 80 observations of each from 4 individual tomograms, were consistent with measurements obtained using cryoEM, cryoET, and dSTORM^37, 38, 51^.

The main drawback of cryoSXT for the analysis of biological samples is the limited resolution of this technique when compared with TEM, cryoEM or cryoET. Whereas TEM and cryoET can reach near-atomic and atomic resolution, respectively^72, 73^, cryoSXT imaging of cells with a 25 nm zone plate can only achieve an effective resolution of approximately 30 nm^32^. This allows imaging of cellular compartments and virus particles, as we demonstrate in this study, but it does not allow the reliable imaging of cytoskeletal components or of individual macromolecular complexes such as proteasomes or ribosomes^75, 76^. The zone plate installed on an X-ray microscope is often outside the control of the end user, but our experience in this study was that use of the 25 nm zone plate did not provide significant additional *biological* information when compared with data collected with the 40 nm zone plate. An additional drawback of cryoSXT is that the relatively low resolution of the images can complicate the differentiation of cellular membrane-bound structures such as autophagosomes, vesicles and other organelles. However, this limitation can be addressed by correlative cryo-microscopy of vitrified samples expressing fluorescent marker proteins, for example the fluorescently-tagged HSV-1 envelope glycoprotein gC that was used in this study to identify sites of virus assembly. Extending the resolution of correlative fluorescence cryo-microscopy using cryoSIM^32^, or increasing the contrast in cryoSXT of specific features in cells via live-cell labelling with metals^77^, are promising future avenues that will address some of the limitations of cryoSXT and extend its utility for biological imaging.

HFF-hTERT and U2OS cells are commonly used for the study of HSV-1 infection^16, 53, 54^. We had originally intended to use only HFF-hTERT cells for this study, to allow comparison with super-resolution fluorescence microscopy studies^16^, but found infected HFF-hTERT cells to be less amenable to interrogation by cryoSXT than other cell lines, such as U2OS cells. We therefore explored the differences in the dynamics of viral infection between HFF-hTERT and U2OS cells using the timestamp virus. In general, the modifications to cellular compartments observed in this study largely replicated those observed in HFF-hTERT cells^16^, suggesting the interactions between viral components and cellular compartments are broadly similar in these two cell types. We observed subtle differences between the infections in these cells, including a change in the nuclear-to-cytoplasmic translocation of the early viral protein ICP0 (Fig. 2). Residues important for the nuclear import/export dynamics of ICP0 have previously been identified: ICP0 possesses a canonical nuclear localization signal at residues 500–506 and deletion of 57 residues from the C terminus abolishes nuclear export of ICP0^78^. Although residues important for trafficking of ICP0 have been mapped, the cellular proteins involved in ICP0 trafficking have yet to be identified. In this study, a higher intensity of eYFP-ICP0 was detected in the nucleus compared with the cytoplasm of infected U2OS cells at every timepoint. In contrast, higher cytoplasmic intensity of ICP0 is observed at late stages of infection in HFF-hTERT cells and other cell lines^16, 79, 80^. This suggests that the expression of host proteins that regulate nuclear import and/or export of ICP0 may differ in U2OS cells. Several host proteins are known to participate in the nuclear trafficking of EP0, the pseudorabies virus orthologue of ICP0: Ran, Importin α1, α3, α7, β1, and transportin-1^81^. Future work is required to identify whether U2OS cells are depleted or enriched in proteins involved in nuclear import/export of ICP0, which may illuminate the mechanisms regulating subcellular localisation of this important viral E3 ligase during infection.

Compared with uninfected U2OS cells, infected cells had a greater local concentration of detectable vesicles in the juxtanuclear space (Fig. 4), consistent with previous research into the distribution of vesicles during HSV-1 infection^16^. For instance, early endosomes and lysosomes have been shown to accumulate at the juxtanuclear region during HSV-1 infection of HFF-hTERT cells^16^. This reorganization of vesicle distribution may be related to a change in microtubule dynamics during infection. Previous studies of HFF and Vero cells have shown that γ-tubulin and pericentrin, which are components of the MTOC, become dispersed during alphaherpesvirus infection, suggesting breakdown of the MTOC^25^. Thereafter microtubules polymerize at multiple foci in the cytoplasm rather than at a single site and the growth rate, length, and stability of nascent microtubules become reduced compared with uninfected cells^25^. We observed a decrease in the abundance of long microtubule filaments and a disappearance of the MTOC as the infection progressed in U2OS cells (Fig. 8), consistent with these previous studies. As the morphology of microtubules changes, the transport of vesicles towards the cell periphery may become obstructed. This may result in the accumulation of vesicles at juxtanuclear regions and may partly explain the increased local concentration of vesicles we observed.

An additional source of new vesicles may arise from the fragmentation of the Golgi complex during HSV-1 infection^64^. Most of the evidence for Golgi fragmentation is based on the dispersion of several Golgi markers (β-COP, Giantin, GM130, 58K protein, and beta-1,4-galactosyltransferase 1) throughout the cytoplasm during HSV-1 infection as assessed by fluorescence microscopy^16, 17, 64^. Golgi fragmentation has been studied to a lesser extent by ultrathin section EM, revealing that cisternae become swollen and separated during infection^17^. Golgi fragmentation is thought to be a consequence of disrupted microtubule dynamics and can be induced by treatment with nocodazole, an inhibitor of β-tubulin polymerization^64^. Although our results are consistent with these observations, the lack of markers for different types of vesicles meant that we could not determine if the vesicles we observed with cryoSXT were Golgi-derived, of endosomal origin, or were unrelated to these cellular compartments. We observed a reduction in the mean size of vesicles as the infection progressed (Fig. 4B), which could arise either from fragmentation of the Golgi complex into small vesicles or an inability of small vesicles to be trafficked from the juxtanuclear region to their target organelles via microtubule transport. Furthermore, using SIM super-resolution fluorescence microscopy we observed infection-associated fragmentation of membranes labelled with the *cis*-Golgi and *trans*-Golgi network components GM130 and TGN46, respectively, in U2OS cells (Fig. 6 and 7). It would be interesting in the future to use fluorescent markers and correlative cryoSIM plus cryoSXT imaging to identify precisely which cellular compartments are found with an increased concentration at the juxtanuclear region of HSV-1 infected cells^32, 82^.

In addition to an increase in the number of vesicles, we observed a significant increase in the median size of lipid droplets during early but not late stages of infection when compared to uninfected cells (Fig. 4C). This observation is similar to a recent study that demonstrated EGFR-mediated upregulation of lipid droplets early in HSV-1 infection (8 hpi) and an increase in lipid droplet size when cells were exposed to dsDNA^83^. Furthermore, the authors of this study demonstrated that accumulation of lipid droplets is transient, returning to baseline within 72 hours following stimulation of cells with dsDNA^83^. While we also observe a transient increase in lipid droplet size following HSV-1 infection, we did not observe a striking increase in the number of lipid droplets per cryoSXT tomogram. However, we note that in our study we infected U2OS cells whereas the previous work used HSV-1–infected primary astrocytes. The authors observed that the increase in size of lipid droplets upon stimulation with dsDNA is cell-type specific, with no increase being observed in THP-1 cells; it is similarly possible that U2OS cells could have a larger number of lipid droplets in the resting state such that the absolute abundance of lipid droplets is not increased in response to HSV-1 infection. An increase in lipid droplet size has also been observed for human cytomegalovirus, a related herpesvirus, after infection for 1–4 days^84, 85^. Lipid droplets are important cell signalling platforms that have been shown to modulate the anti-viral immune response during infection^83^. The high resolution of cryoSXT when compared with confocal microscopy, combined with the high contrast afforded by carbon-rich lipid droplets, makes cryoSXT imaging particularly suitable for future research into the link between lipid droplet size and cellular innate immune responses.

We observed that mitochondria became more elongated and branched as infection progresses, in some cases forming extensive networks (Fig. 5). Branching of mitochondria can either occur via *de novo* synthesis or by fusion of mitochondria^24, 86^, and there are several possible explanations for the change in mitochondrial morphology observed during HSV-1 infection. Mitochondrial movement tends to occur along microtubules and this movement influences mitochondrial fusion/fission dynamics. Fission can arise from divergent movement of mitochondrial extensions along microtubules and fusion is supported by convergent movement of mitochondria^87^. Nocodazole treatment to depolymerize microtubules blocks transport, fusion and fission of mitochondria, and there is evidence that thin microtubule extensions develop when fission is obstructed^88^. It is possible that fission of existing mitochondrial networks may be obstructed when microtubules depolymerize during HSV-1 infection, and this may prevent the generation of small mitochondria. Such changes to the microtubule network begin at 6 hpi and would thus be expected to have a greater influence on mitochondrial morphology in the late stages of infection^25^, consistent with our observations. Alternatively, the morphological changes to mitochondria may reflect a cellular response to increased respiratory demand^89^. An increase in ATP production can be achieved by mitochondrial elongation, for example under conditions of stress such as hypoxia and starvation of glucose metabolism^19, 20^. An increase in respiration, including oxidative phosphorylation, has been observed during human cytomegalovirus infection^90^. An increased number of elongated mitochondria in cells at late stages of infection could increase ATP production during infection. Increased oxidative stress provides a third plausible explanation for the observed changes in mitochondrial morphology. Production of reactive oxygen species (ROS) during respiration appears to be a common feature of viral infection that has been observed for hepatitis C virus, respiratory syncytial virus and the herpesvirus Epstein-Barr virus^21–23^. One mechanism by which the cell responds to oxidative stress is by fusion of undamaged and ROS-damaged mitochondria to allow for compensatory effects by sharing resources needed for ATP production^24^. It is possible that changes in mitochondrial morphology we observed may have arisen via enhanced fusion in response to increased oxidative stress during infection.

Although a change in energy metabolism may reflect a generalized response by the cell to infection, mitochondrial elongation has been observed during infection with other viruses (such as dengue virus) that inhibit mitochondrial fission^91^. Several HSV-1 proteins have been reported to localize at mitochondria (pUL7, pUL16, pUS3, pUL12.5), suggesting that HSV-1 directly modulates mitochondrial activity^92–95^. pUS3 inhibits the activity of electron transport chain complexes II and III as early as 6 hpi^93^ and pUL12.5 functions in the depletion of mitochondrial DNA and downregulation of mitochondrial proteins, including ND6 and COX2, as early as 4–8 hpi^94^. The functional consequences of pUL16 binding mitochondria are not well characterized, although we note that a pUL16 mutant co-localizes with mitochondrial fission sites^95^. The precise mechanisms by which HSV-1 alters the architecture of mitochondria and the role of specific viral proteins, versus virus-induced metabolic strain, thus remains unclear. Combining metabolic profiling of infected cells with ultrastructural analysis of mitochondrial morphology, using wild-type and mutant (knock-out) viruses, will help illuminate the factors that drive the dramatic remodelling of mitochondria observed during HSV-1 infection and the functional consequences thereof.

In conclusion, we have demonstrated that cryoSXT produces quantitative high-resolution 3D data for biological research by studying the ultrastructural changes to cellular compartments induced during HSV-1 infection. CryoSXT allows the detection of HSV-1 capsids and virions in distinct subcellular locations, such as the nucleus, perinuclear space, cytoplasmic vesicles, and cell surface. Use of the timestamp HSV-1 reporter virus facilitated identification of individual cells at early or late stages of infection. In these subpopulations we observed accumulation of vesicles at juxtanuclear assembly compartments, a transient increase in the size of lipid droplets, and elongation plus branching of mitochondria as infection progresses. The ability of cryoSXT to image the entire depth of infected cells in a near-native state, with minimal sample processing, highlights its utility as a tool for 3D imaging to identify changes in cellular architecture that accompany virus infection.

## Materials & Methods

### Reagents

250 nm gold colloid fiducials were purchased from BBI Solutions (EM.GC250, batch 026935). The working mixture was prepared via sedimentation of 1 mL of stock solution by centrifugation (12×g, 5 mins, RT) and then resuspending the pellet in 50 µL Hanks’ Balanced Salt Solution (HBSS; Thermo Fisher). The fiducials were sonicated at 80 kHz (100% power) and 6°C to prevent aggregation. 3 mm gold EM finder grids with a perforated carbon film (R 2/2, 200 mesh) were purchased from Quantifoil (AU G200F1 finder, batches Q45352 & Q45353). Poly-l-lysine was purchased from Sigma Aldrich.

### Cell Lines

Mycoplasma-free U2OS cells (ATCC HTB-96; RRID CVCL_0042) and human foreskin fibroblast cells immortalized with human telomerase reverse transcriptase (HFF-hTERT cells)^96^ were cultured in Dulbecco’s Modified Eagle’s Medium (DMEM; Thermo Fisher) supplemented with 10% (v/v) foetal bovine serum (FBS; Capricorn), 2 mM l-glutamine (Thermo Fisher), and 100 U/mL penicillin/streptomycin (Thermo Fisher). HBSS and 0.25% Trypsin-EDTA (Thermo Fisher) were used to wash and detach adherent cells, respectively. Cells were maintained in a humidified 5% CO_2_ atmosphere at 37°C.

### Biosafety Measures

All cells and viruses were handled according to containment level 2 (CL2) guidelines and a risk assessment was carried out and approved by the Rutherford Appleton Laboratory (RAL) Health and Safety Committee. EM grids containing cells and viruses were handled in appropriate microbiology safety cabinets and forceps were regularly washed in 70% (v/v) ethanol. Personal protective equipment in the form of lab coats, lab gloves, and goggles were worn to protect experimenters. All tissue-culture, cryocooling, and imaging equipment was stored in CL2 laboratories.

### Recombinant Viruses

Infections were performed using recombinant HSV-1 strain KOS expressing either the endogenously tagged viral proteins eYFP-VP26 and gM-mCherry (Fig. 1) or the endogenously tagged viral proteins eYFP-ICP0 and gC-mCherry (timestamp HSV-1, Fig. 2–8 and Supp. Fig. 1–4)^16^, to allow distinction between early and late stages of infection in U2OS and HFF-hTERT cells, with the exception of the leftmost panel in Fig. 1I for which a non-fluorescent wild-type HSV-1 strain KOS was used. Virus stocks were prepared by infection of Vero cells at low MOI (0.01) for 3–5 days, until cytopathic effect was evident, before scraping cells into the medium. The cells were frozen at −70°C, thawed and sonicated at 50% amplitude for 40 seconds. Crude virus stocks were clarified by centrifugation at 3,200×g in a benchtop centrifuge, aliquoted, and viral titers of the aliquots were quantified on Vero and U2OS cells as described previously^97^.

### Infection Assays

For widefield imaging under cryogenic conditions and cryoSXT, EM grids were glow discharged and treated with filtered poly-l-lysine for 10 minutes as described previously^33^. 3×10^5^ U2OS or HFF-hTERT cells per well were seeded in 6-well plates containing the treated EM grids and were incubated overnight. Subsequently, the cells were infected with timestamp HSV-1 at an MOI of 1–3. For widefield microscopy of timestamp HSV-1 to measure the progression of replication over time, U2OS and HFF-hTERT cells were allowed to grow overnight following seeding in 6-well plates at 2×10^5^ cells per well (Fig. 2A) or on borosilicate coverslips in 12-well plates at 1×10^5^ cells per well (Fig. 2B). The cells were infected with the recombinant HSV-1 with an MOI of 1–3. For SIM and confocal microscopy, U2OS cells were seeded on borosilicate coverslips in 12-well plates overnight at 1×10^5^ cells per well. The cells were infected with the recombinant HSV-1 with an MOI of 3. The time of inoculation was designated the start time of infection. For all infections, to maximize adsorption of virus, cells were incubated in a low volume of medium (250 µL/well in 12-well plates and 500 µL/well in 6-well plates) for 1 hour in a humidified 5% CO_2_ atmosphere at 37°C and the plates were swirled every 15 minutes. For widefield imaging under cryogenic conditions and cryoSXT, the medium was topped up to 2 mL and the samples were incubated for 9 hours alongside uninfected controls, except for the samples in Fig. 1, which were incubated for 16 hours. The EM grids were overlayed with 2 µL of the gold fiducial working mixture as described in the *Reagents* section. A Leica EM GP2 plunge freezer was used to blot the grids for 0.5– 1 s at 30°C and 80% humidity. The grids were then plunged into liquid nitrogen-cooled liquid ethane and transferred into liquid nitrogen storage before imaging. For the timestamp HSV-1 images (Fig. 2A), the inoculum was diluted to 2 mL with medium for the remainder of the incubation (9–24 hours). For the experiments measuring the progression of timestamp HSV-1 replication over time (Fig. 2B), media was aspirated after the 1 hour of incubation and the cells were treated for 1 min with citric acid (40 mM citric acid pH 3, 135 mM NaCl, 10 mM KCl) to inactivate unabsorbed virus. Cells were then washed thrice with PBS and then overlain with 500 µL of fresh medium before incubation for a further 2–24 hours. For confocal microscopy and SIM, the media were topped up to 1 mL for the remainder of the incubation (see figure legends for varied time points). For cells stained with MitoTracker Deep Red FM (Thermo Fisher), the media was aspirated 30 minutes before fixation and washed twice with serum-free media. 50 nM MitoTracker in serum-free media was added to the cells for 30 minutes. For all samples not prepared on EM grids, the cells were washed twice with HBSS or PBS and were fixed with 4% (v/v) formaldehyde for 20 minutes, followed by three HBSS/PBS washes. For confocal microscopy and SIM, the cells were washed twice with PBS, permeabilised with 0.1% saponin in PBS for 30 minutes at room temperature on a rocking platform and were blocked for 30 minutes with a PBS solution of 0.1% saponin and 5% (v/v) FBS at room temperature on a rocking platform. The samples were stained with either 2.5 μg/mL mouse anti-human GM130 (Clone 35/GM130 (RUO), BD Biosciences, RRID: AB_398142), 10 μg/mL mouse anti-human TGN46 (SAB4200355, Merck), or a 1 in 20 dilution of rat anti-β-tubulin (clone YL1/2) hybridoma supernatant^98^ in blocking solution, washed, then stained with 4 µg/mL goat anti-mouse Alexa Fluor 647 antibody (A-21235, ThermoFisher) in blocking solution, washed and mounted using ProLong Gold containing DAPI (P36930, ThermoFisher).

### Widefield Microscopy

For room temperature samples, a Zeiss AxioImager2 microscope with an achromatic 50× air objective (Zeiss LD EC Epiplan-Neofluar 50x/0.55 DIC M27; NA=0.55) was used to image fixed infected cells grown on plastic 6-well plates. Fluorescent images were collected using the Zeiss 46 HE YFP filter (Excitation 500±25 nm, Emission 535±30 nm) and the Zeiss 64 HE mPlum filter (Excitation 587±25 nm, Emission 647±70 nm). For cryo-widefield microscopy, cells at early- and late-stages of infection were identified based on the spatiotemporal expression of eYFP-ICP0 and gC-mCherry using a Zeiss AxioImager2 microscope with an achromatic 50× objective as described above. A liquid nitrogen cryostage (Linkam Scientific) was used to maintain the samples at 77 K. Each grid was mapped in its entirety in the brightfield and fluorescent channels (as above) using LINK (Linkam Scientific).

### Cryo-Soft-X-Ray Tomography

X-ray images were collected using an UltraXRM-S/L220c X-ray microscope (Carl Zeiss X-ray Microscopy) at beamline B24 at the UK synchrotron Diamond Light Source. Grids were imaged in a liquid nitrogen-cooled vacuum chamber and samples were illuminated with 500 eV X-rays (λ = 2.48 nm) for 0.5 or 1 s per projection. The transmitted light was focused by diffraction using zone plate objectives with nominal resolution limits of either 25 nm or 40 nm. The 25 nm zone plate offers higher resolution but captures a smaller field of view (∼10×10 μm) than the 40 nm zone plate (∼16×16 μm). Only one zone plate can be installed in the microscope and the zone plate is not user changeable. The installed zone plate differed across the beam time allocations use for this study, with all images being collected using the 25 nm zone plate except for Fig. 1 C, D, **and** G–I, for which the 40 nm zone plate was used. Transmitted images were collected using a 1024B Pixis CCD camera (Princeton instruments). X-ray mosaic images (7×7 images capturing 66.2×66.2 µm for the 25 nm objective and 106.0×106.0 µm for the 40 nm objective) were collected from different areas on the grid to assess overall cell morphology. For identification of early and late stages of infection, X-ray mosaics were compared with fluorescent scans acquired on the cryo-widefield microscope to identify specific infected cells. These mosaics were also used to identify regions of interest for tomography. Tilt series of projections were collected from these regions by rotating the sample around an axis normal to the incident X-ray beam by up to 140° in increments of 0.2° or 0.5° per image, with maximum tilt angles of −60°/+60° and −70°/+70° for the 25nm and 40nm objective, respectively. SXT tilt series were processed using IMOD (version 4.9.2)^42^. The images were aligned along a single axis. A coarse alignment was performed by cross-correlation with a high frequency cut-off radius of 0.1. Coarsely aligned tilt series were further aligned manually using gold fiducials and dark cellular compartments, such as lipid droplets. A boundary model was generated to reorient the 3D data in case the sample was collected at an angle and final alignment was performed using linear interpolation. Tomograms were generated using back projection followed by 20 iterations of a simultaneous iterations reconstruction technique (SIRT)-like filter to reduce noise.

### Structured Illumination Microscopy and Deconvolution

A custom, three-colour SIM microscope^99^ was used to collect images. A ferroelectric binary Spatial Light Modulator (SLM) (SXGA-3DM, Forth Dimension Displays) was used to pattern the light with a grating structure (3 angles and 3 phases). Light was collected by a 60× water immersion objective with a NA of 1.2 (UPLSAPO60XW, Olympus) and a sCMOS camera (C11440, Hamamatsu). eYFP-ICP0 fluorescent emission was captured using a 488 nm laser (iBEAM-SMART-488, Toptica) and an BA510-550 (Olympus) emission filter. gC-mCherry fluorescence was captured using a 561 nm laser (OBIS561, Coherent) and a BrightLineFF01-600/37 filter (Semrock). AF647 fluorescence was captured using a 640 nm laser (MLD640, Cobolt) and a BrightLineFF01-676/29 filter (Semrock). Background-reduced and resolution-enhanced images were reconstructed from raw SIM data using FairSIM^100^. Deconvolution was performed alongside using a Richardson-Lucy algorithm with 5 iterations. 100 nm beads (TetraSpeck Microspheres, Thermo Fisher) were used to determine the shifts in X and Y for channel alignment. The channels were aligned using the TransformJ plugin in Fiji^101^.

### Confocal microscopy

A Zeiss LSM700 inverted confocal AxioObserver.Z1 microscope was used at room temperature to capture images with ZEN software (Zeiss). A 100× apochromatic objective with oil immersion and pinhole set to 1 airy unit was used to collect images. Z stacks containing 1024×1024 pixels at 400 nm increments were captured within a 16-bit unsigned range using 8-fold line averaging. Maximum Z projections were generated using Fiji^101^.

### Segmentation

Mitochondria were segmented using *Contour*, a bespoke semi-automated segmentation and quantitation tool developed with Python 3 (full details on *Contour* can be found in reference 60). Briefly, *Contour* automatically segments high contrast features such as mitochondria by thresholding and then applying a restriction on the minimum number of consecutive segmented pixels vertically and horizontally. Next, gaps in the segmented volume can be filled in by running this algorithm in local regions of interest. Separate elements in the segmented volume are differentiated by grouping of neighbouring voxels together. The differentiated elements are colour-coded and their volumes are quantitated from the number of voxels. The edges of the segmented elements are smoothened in each image plane by translating the image by one pixel in all eight cardinal and ordinal directions in the XY plane and calculating the median pixel value for all these translations. A 3D Gaussian filter with a sigma of 2 was also added using Fiji to further smoothen the elements^101^. In *Contour*, the width of each segmented element was calculated by finding all the coordinates of voxels at the perimeter of segmented elements and calculating the largest modulus between any two coordinates. Segmented volumes of cytoplasmic vesicles were generated manually using the Segmentation Editor 3.0.3 ImageJ plugin^101^ and these were imported into *Contour* to differentiate between segmented elements and quantitate the width of the vesicles. Segmented volumes were visualized in 3D using the 3D Viewer plugin in ImageJ^101^. Cytoplasmic vesicles were segmented from 12 well-reconstructed X-ray tomograms lacking X-ray damage. Lipid droplets were automatically segmented from 94 tomograms using *Contour* from 8-bit tomograms. Tomograms were excluded from segmentation if they were poorly reconstructed, were subjected to X-ray damage, or were from thick sections and contained out-of-focus lipid droplets in some projection planes. Given that gold fiducials and lipid droplets have similar projection intensities in 8-bit images, gold fiducials were also included in the automatic segmentation. During curation of the segmented volumes, gold fiducials were manually erased. This was possible because they are easy to distinguish from lipid droplets based on the higher intensity of their missing wedge artefacts. The projection intensities of lipid droplets and gold fiducials are lower than other material in the tomograms, including noise, and they could thus be segmented based purely on threshold values without applying a width restriction. This prevented the exclusion of very small lipid droplets. Volumes were calculated in *Contour* for each lipid droplet. Lipid droplets that could not be individually resolved or were cut off at the edges of tomograms were excluded from the analysis.

### Graphs and statistics

Distributions of capsid and virion widths were illustrated using a Violin SuperPlot^102^, with data grouped by source tomograms. The stacked area plots for the proportion of infected cells at different stages of infection were generated using the ggplot2 package^103^ in R studio^104^. The distribution of vesicle widths were illustrated using a SuperPlot^105^, with data grouped by source tomograms. The numbers of mitochondrial branch points (branching nodes) were illustrated using a Violin SuperPlot^102^, with data grouped by replicate. A two-tailed paired t-test was used to compare the width of the nuclear envelope at a site of primary envelopment with the width of the nuclear envelope elsewhere on the same tomogram using Excel (Microsoft). The t-test was two-tailed because we observed a normal distribution of widths and it was paired because the two data points (width of nuclear envelope at perinuclear viral particle and in nearby region) were collected from the same tomogram. The variance in the width of the extracellular virions was greater than four times the variance in the width of nuclear capsids. As a result, a Mann-Whitney *U* test for unequal variance was used to assess the significance of the difference in width of capsids and virions using R Studio^104^. To assess significance of differences in mean vesicle widths, a one-way ANOVA and Tukey tests were performed using Prism version 8.2.1 (GraphPad Software). These tests were used instead of t tests to avoid the higher risk of type I errors associated with performing multiple t tests on more than two conditions. For the same reason, one-way ANOVA and Tukey tests were used to assess significant differences in the number of mitochondrial branching nodes (using R Studio^104^). The data distributions for the lipid droplet volumes were positively skewed and median volumes were chosen for further analysis because they are less affected by extreme values than means. Owing to the skew, Mann-Whitney *U* tests were used to determine significance of differences between conditions for the lipid droplet volumes. The tomograms used to quantitate the number of branching nodes were selected according to the following criteria. Firstly, we excluded tomograms that were poorly reconstructed (e.g. due to displacement of the rotational axis during data collection or due to a lack of fiducials or lipid droplets for image alignment). Next, we only included tomograms in which the mitochondria were well dispersed throughout the field of view to ensure that we didn’t systematically underestimate the number of branching nodes by including tomograms where only small fragments of individual mitochondria were visible, for example in the corners of the field of view. Finally, we excluded tomograms that contained swollen mitochondria because this was taken to indicate that the cells might be undergoing apoptosis, a process known to cause significant alteration to mitochondrial morphology^61^. We assessed an equal number of tomograms for each replicate and condition to avoid introducing unequal variance into our ANOVA test.

## Acknowledgements

We thank Diamond Light Source for access to beamline B24 (via response mode applications mx18925, mx19958, bi21485 and bi23508) and the experimental hall coordinators for helpful support. We thank members of beamline B24 at the Diamond Light Source (Mohamed Koronfel, Ilias Kounatidis, Chidinma Okalo, and Matt Spink) for technical support with cryoSXT. We thank João Ferreira Fernandes (University of Oxford) and Thomas Fish (Diamond Light Source) for help with the development of *Contour*. We thank David Carpentier (University of Cambridge) for technical support with confocal microscopy. We thank Harriet Groom (University of Cambridge) for reading and advising on the manuscript. This work was supported by a PhD studentship co-funded by Diamond Light Source and the Department of Pathology, University of Cambridge, to KLN, by a research fellowship from the Deutsche Forschungsgemeinschaft to KMS (DFG; SCHE 1672/2-1), by the funding from the Engineering and Physical Sciences Research Council (EPSRC; EP/H018301/1) and the Medical Research Council (MRC; MR/K015850/1 and MR/K02292X/1) to CFK, by a Biotechnology and Biological Sciences Research Council (BBSRC) Research Grant (BB/M021424/1) to CMC, and by a Sir Henry Dale Fellowship, jointly funded by the Wellcome Trust and the Royal Society (098406/Z/12/B) to SCG. For the purpose of Open Access, the authors have applied a CC BY public copyright licence to any Author Accepted Manuscript (AAM) version arising from this submission.

## Data Availability

Original imaging data for tomograms illustrated in the manuscript are deposited with the University of Cambridge Apollo Repository, available at https://doi.org/10.17863/CAM.78593. Representative tomograms have also been published in the EMPIAR repository (EMBL-EBI) with the accession number EMPIAR-10972.

---

**Supp. Fig. 1.**
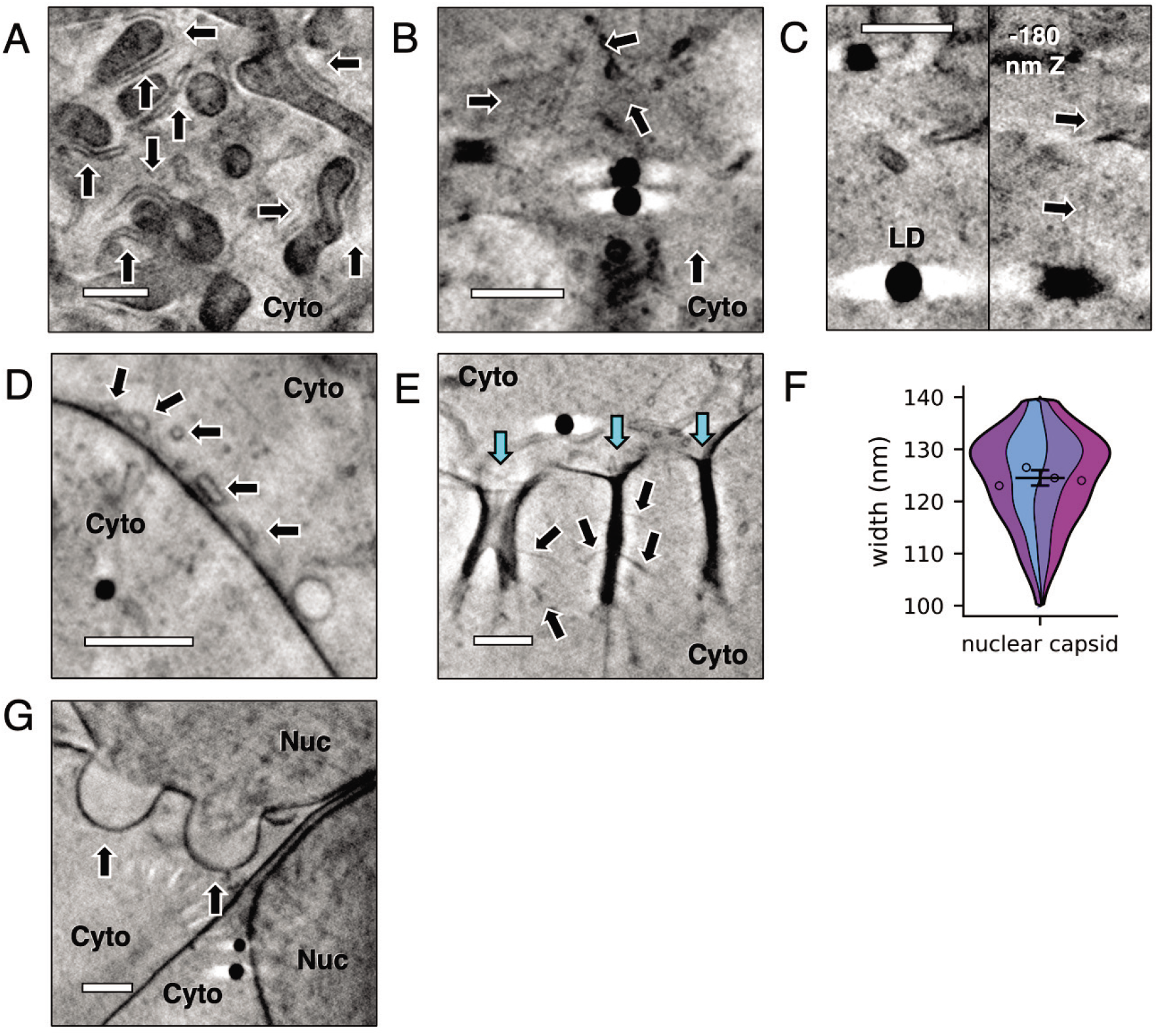
High resolution structures visible with the 25 nm zone plate objective. 139 CryoSXT tomograms were recorded from 107 cells using a 25 nm zone plate objective and several structures that were unrelated to HSV-1 infection were observed, including some that were not visible using the 40 nm zone plate objective. (**A**) The endoplasmic reticulum (ER) forms a silhouette (arrows) around the mitochondria and the ER lumen is visible with the 25 nm zone plate. Cyto, cytoplasm. (**B**) Linear structures resembling cytoskeletal filaments are visible with the 25 nm zone plate (arrows). (**C**) A putative cytoskeletal filament (arrows) is in close apposition to a lipid droplet (LD) and may represent a physical interaction. (**D**) Small vesicles with widths of 150–300 nm in the peripheral cytoplasm are observed (arrows). (**E**) Large internalisations of the plasma membrane with depths of 1.6–2.2 µm (cyan arrows) and smaller side extensions (black arrows) are visible and may represent events of clathrin-independent bulk endocytosis^108^. (**F**) The width of nuclear capsids was remeasured after imaging with the 25 nm zone plate: 124.5 nm ± 0.96 nm SEM (n=80 from 4 tomograms; 8.55 nm SD). (**G**) Bulging of the nuclear envelope is observed (arrows). We initially observed these in HSV-1 infection and thought it may represent a virus-directed decrease in the integrity of the nuclear envelope, but we found multiple examples in uninfected cells suggesting that they are a characteristic of U2OS cells. Nuc, nucleus. Scale bars = 1μm.

**Supp. Fig. 2.**
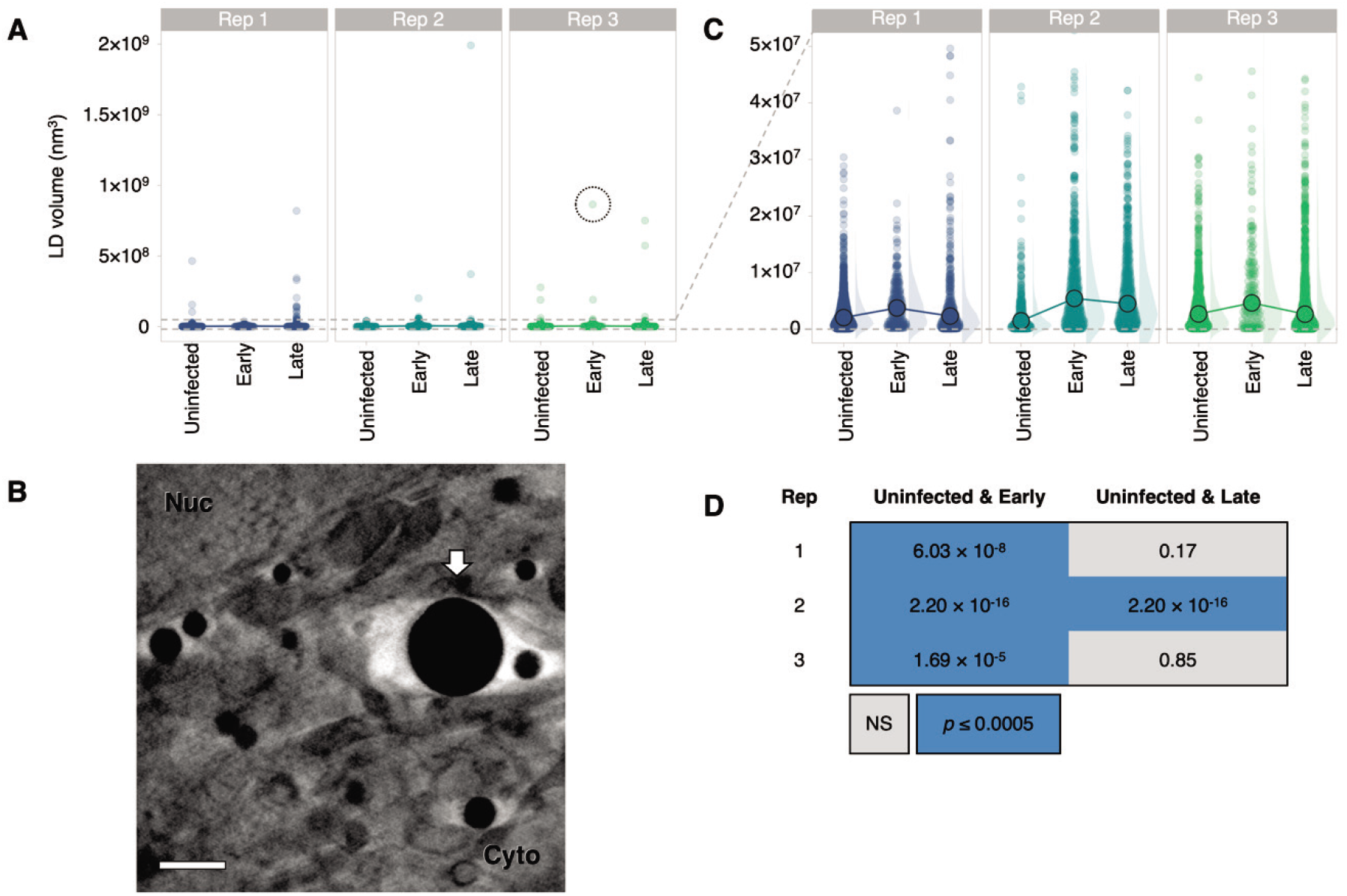
Lipid droplets transiently increase in size during early stages of HSV-1 infection. 4845 lipid droplets across the three replicates were segmented using *Contour*^60^ and their volumes were calculated. Scale bars = 1 µm. (**A**) A linear plot of lipid droplet volumes reveals a similar number of extremely large lipid droplets (> 5×10^7^ nm^3^) in U2OS cells. The circled lipid droplet is shown in (**B**). (**B**) A large lipid droplet observed in a U2OS cell at an early stage of infection. Scale bar = 1 µm. (**C**) A linear plot of lipid droplet volumes, truncated at 5×10^7^ nm^3^, reveals that in all conditions lipid droplet volumes were positively skewed rather than normally distributed. Median volumes are shown (large circles) because they are less affected by extreme values than mean volumes. The median lipid droplet volume was highest in cells at early stages of infection for all three replicates. (**D**) Given that the distributions were positively skewed, non-parametric Mann-Whitney *U* tests were carried out to determine significant differences at a 0.0005 *p-*value threshold. NS, no significant difference. In all three biological replicates the lipid droplets of cells at an early stage of infection are larger than in uninfected cells. The lipid droplets in cells at late stages of infection are not significantly larger than in uninfected cells for two of the three biological replicates, suggesting that a transient increase in lipid droplet volume accompanies HSV-1 infection of U2OS cells.

**Supp. Fig. 3.**
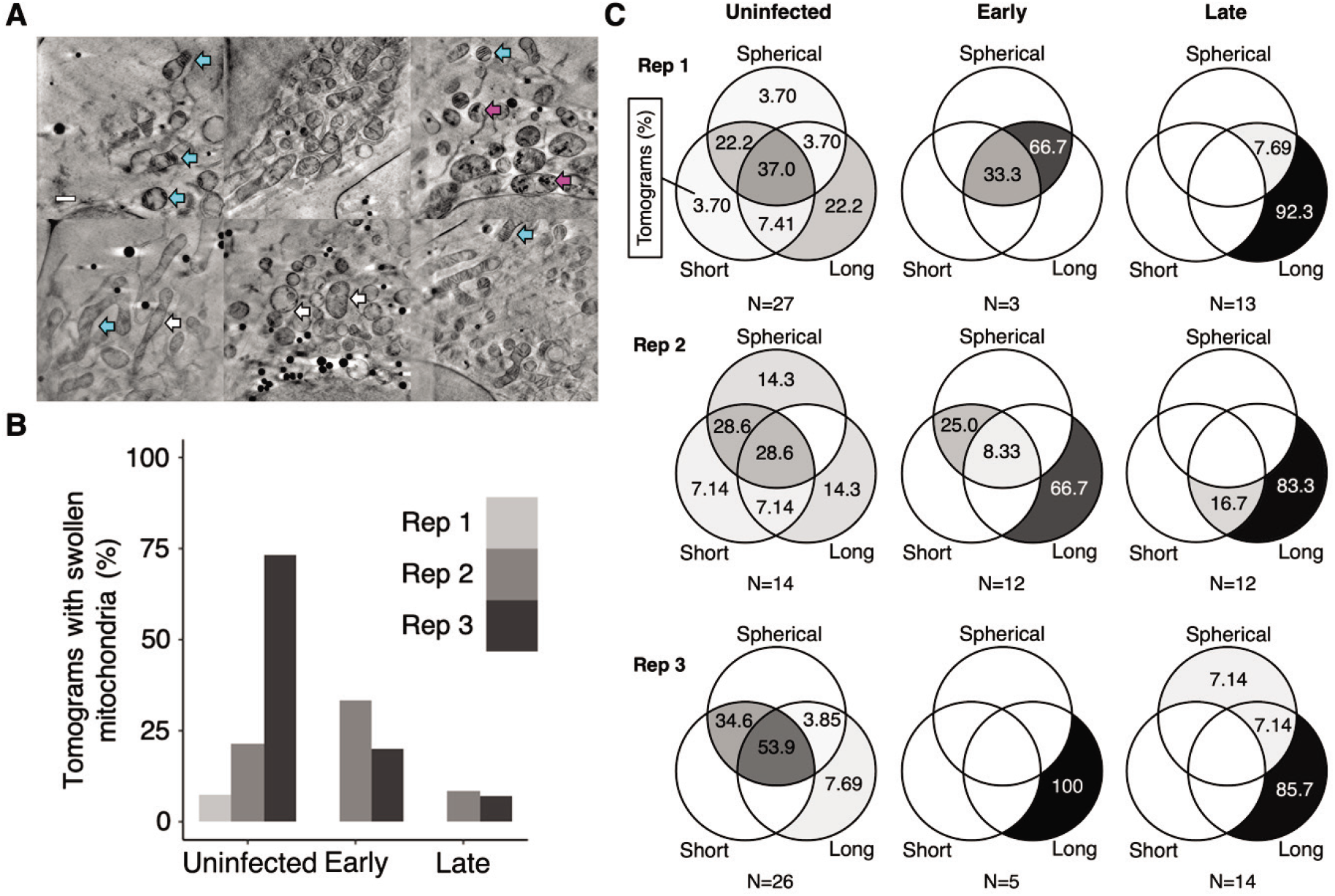
The heterogenous morphology of mitochondria. Heterogenous mitochondrial morphologies are observed in cryoSXT tomograms collected from uninfected cells and cells at early and late stages of infection with timestamp HSV-1. Scale bars = 1 µm. (**A**) In some cases, mitochondria have light matrices with highly contrasting cristae (cyan arrows). This “swollen” phenotype has been reported to occur during cytochrome *c* release from porous mitochondria during apoptosis^61^. Dark matter is also observed in the matrix (magenta arrows) and may represent vesiculation. Small dark puncta are present in the matrix (white arrows) and could represent vesicles or short cristae. (**B**) The percentages of tomograms with swollen mitochondria for uninfected cells and cells at early- or late-stages of infection in three independent replicates. (**C**) The percentages of tomograms collected from uninfected cells and those at early- or late-stages on infection in each replicate that contain different combinations of mitochondrial morphologies.

**Supp. Fig. 4.**
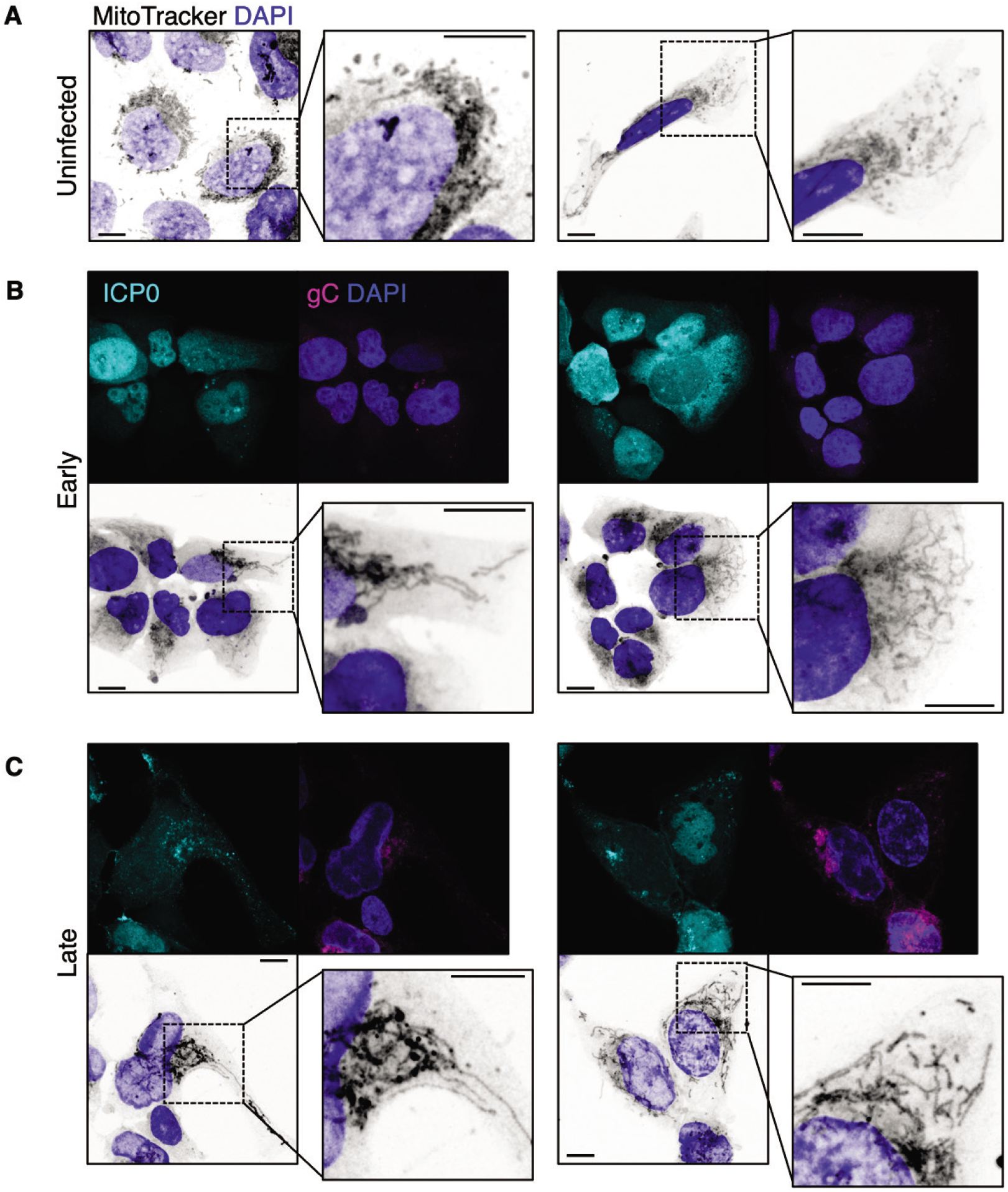
Confocal imaging of mitochondrial morphology. U2OS cells infected with timestamp HSV-1 (MOI 3) were fixed at indicated times and imaged by confocal microscopy. Mitochondria were stained with MitoTracker Deep Red FM. Scale bars = 10 μm. (**A**) Mitochondria in uninfected cells were morphologically heterogenous. (**B, C**) In cells at (**B**) early (6 hpi) and (**C**) late (16 hpi) stages of infection, a greater proportion of elongated mitochondria were observed.

**Supp. Video 1. Segmentation of vesicles and mitochondria in the cytoplasm of a cell at a late stage of infection.** CryoSXT data was collected from U2OS cells infected for 9 hours with the timestamp HSV-1 virus at an MOI of 1. Cryo-fluorescence microscopy revealed that this cell was at a late stage of infection. The mitochondria were segmented using *Contour*^60^ and separate mitochondria are colour-coded in shades of orange, red, pink and purple. Cytoplasmic vesicles were segmented using *Segmentation Editor* in *ImageJ*. The vesicles were later differentiated and color-coded using *Contour*^60^ and are displayed here in shades of blue and green. Field of view is 9.46×9.46 µm.

**Supp. Video 2. Segmentation of cytoplasmic vesicles reveals the effect of HSV-1 infection on vesicle concentration at juxtanuclear sites.** CryoSXT data was collected from uninfected U2OS cells and U2OS cells infected for 9 hours with the timestamp HSV-1 virus at an MOI of 1. Cytoplasmic vesicles were segmented using *Segmentation Editor* in *ImageJ*. The vesicles were later differentiated and colour-coded using *Contour*^60^. Fields of view are 9.46×9.46 µm.

**Supp. Video 3. Segmentation of mitochondria reveals the effect of HSV-1 infection on mitochondrial morphology.** CryoSXT data was collected from uninfected U2OS cells and U2OS cells infected for 9 hours with the timestamp HSV-1 virus at an MOI of 1. Mitochondria were segmented and colour-coded using *Contour*^60^ and appear elongated and branched in cells at late stages of infection. Fields of view are 9.46×9.46 µm.

